# Mitochondria transplantation between living cells

**DOI:** 10.1101/2021.11.09.467932

**Authors:** Christoph G. Gäbelein, Qian Feng, Edin Sarajlic, Tomaso Zambelli, Orane Guillaume-Gentil, Benoît Kornmann, Julia A. Vorholt

## Abstract

Mitochondria and the complex endomembrane system are hallmarks of eukaryotic cells. To date, it has been difficult to manipulate organelle structures within single live cells. We developed a FluidFM-based approach to extract, inject and transplant organelles from and into living cells with subcellular spatial resolution. The approach enabled the transfer of controlled quantities of mitochondria into cells while maintaining their viability and monitoring their fate in new host cells. Transplantation of healthy and drug-impaired mitochondria into primary keratinocytes allowed real-time tracking of mitochondrial subpopulation rescue. Fusion with the mitochondrial network of recipient cells occurred 20 min after transplantation and continued for over 16 hours. After transfer of mitochondria and cell propagation over generations, we show that donor mtDNA was replicated in recipient cells without the need for selection pressure. The approach opens new prospects for the study of organelle physiology and homeostasis, but also for mechanobiology, synthetic biology, and therapy.

## Introduction

Single-cell surgery approaches promise minimally invasive perturbation, i.e. removal or introduction of cellular compartments without compromising cell viability. Manipulation of mitochondria receives special emphasis due to their central cellular role: they are at the heart of energy conversion and link cellular metabolism to signaling pathways and cell fate decision (1–4). Mitochondria harbor their own genetic content (mtDNA), which is prone to accumulating erroneous, disease causing mutations (5–7) and are subject to quality control (8, 9). Although mitochondria are generally inherited strictly vertically to daughter cells, exchange of larger cellular components including mitochondria has also been observed in tissues of multicellular organisms (10–13). To reconstitute such transfer events, therapy approaches involve the grafting of purified mitochondria into a damaged area of a tissue or their intravenous injection (14). However the fate of these mitochondria is unknown (15).

With limited means to study and quantify the transfer of mitochondria into cells and without ways to analyze dose-response relationships experimentally, it is difficult to gain mechanistic insight on the actual impact of cytoplasmic and mitochondrial transmissions under healthy or diseased cellular states. Extraction and injection of organelles from and into single cells is technically demanding. Miniaturized probes have a high potential to manipulate and sample individual cells within their microenvironment at high spatiotemporal resolution (16). Nano-scaled pipettes and nanotweezers allow sampling and trapping of individual charged molecules and single mitochondria (17, 18) and have been combined with –omics methods, enabling compartment-resolved single cell studies (19, 20). Other specialized microfluidic devices for microinjection into cultured cells have been introduced (21). Using a modified microcapillary pipet, mitochondria injection was achieved (22). However, micropipette-based approaches are limited in terms of volume scalability, have only been applied to single mitochondria, and have been limited to either cell extraction or cell injection. In addition, the success rate of transferring mitochondria into single cultured cells has been low and requires use of cells artificially depleted of mtDNA with subsequent selection of transformed cells. This limits the approach to selective conditions and, in particular, it has prevented studies on the dynamic behavior of mitochondrial subpopulations to this point.

Despite all these crucial developments in single-cell technologies, functional transfer, i.e. transplantation of organelles from cell to cell, has not yet been achieved with exception of much larger oocytes.

FluidFM (23) combines the high-precision force-regulated approach of an atomic force microscope (AFM, pN to µN) with the volumetric dispensing of nano-scale pipets (fL to pL) under optical inspection, providing the forces and volume control relevant for single-cell manipulation (24). These features are unique among miniaturized probes and pivotal for driving the probe into the cytosolic compartment in a minimally invasive manner when delivering and extracting molecules (including plasmids, RNAs, and proteins) into- and from viable cells (25, 26). In this study, we established FluidFM as a single-cell technology for intra- and intercellular micromanipulation of organelles in living eukaryotic cells (Fig. 1A-C). The fluidic handling of subcellular compartments poses a challenge for miniaturized probes because endomembrane structures are relatively large (mitochondria: 300-800 nm in diameter (27)) and form interconnected networks. Organelle manipulation also bears a higher risk of rupturing the cytoplasmic membrane compared to the few nm opening of probes used for molecule injection and extraction in prior work (25, 26). Organelle extraction and injection achieved here required dedicated fabrication of tips with customized aperture area (*A*), to both overcome steric constraints and increase the range of applicable suction forces (*F*_*max*_) at the aperture via the fluidic pressure (*Δp*) in accordance to *F*_*max*_ = *Δp* × *A*. We adapted the size and geometry of the aperture and microchannel of the FluidFM probes, and used the tips in combination with force control for membrane insertion via automated AFM and exertion of suction (i.e. tensile) forces via automated pressure controller. We simultaneously inspected target cells and the transplant inside the transparent cantilever (via phase contrast and fluorescence microscopy, Figure 2A-B). We demonstrate the establishment of a robust and maximally efficient method for the extraction of mitochondria from living cells and for functional transplantation between individual cells. The gentle mode of probe insertion and extraction ensures cell viability and allows monitoring of mitochondrial dynamics in real time and across cell generations, and tracking mitochondrial variants and fate in native cells.

**Fig. 1.**
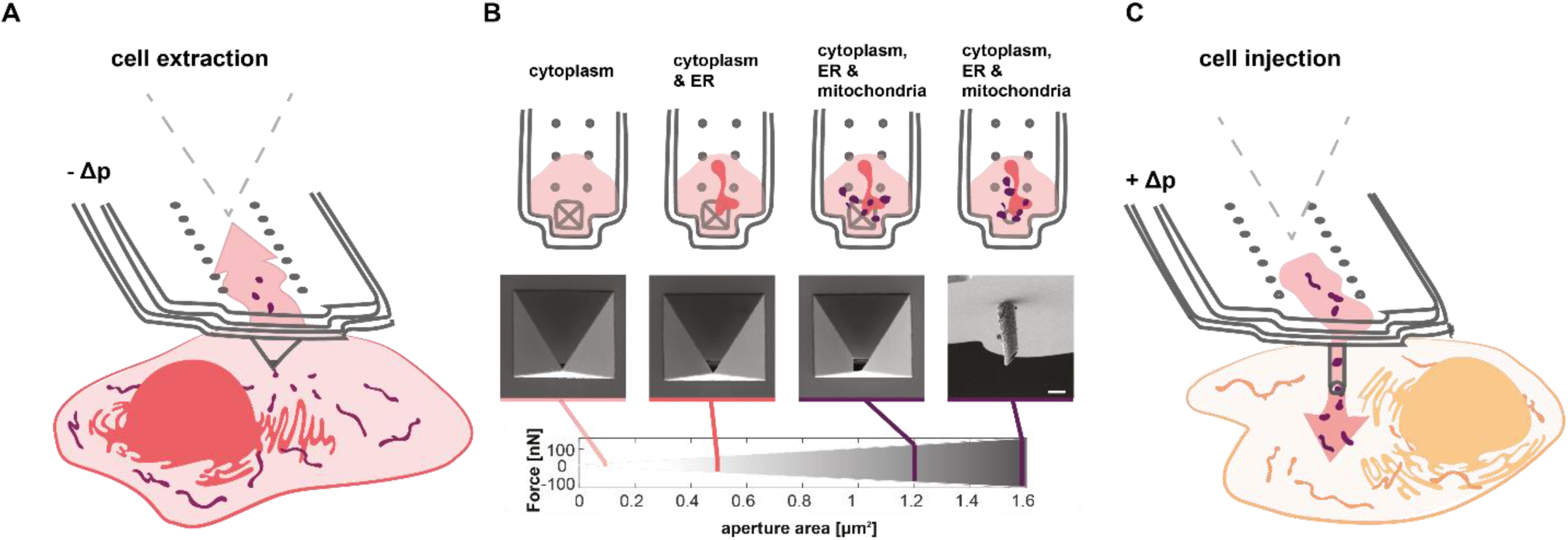
(**A**) Schematic of organelle extraction using FluidFM. Extraction volumes are tuned by applying negative pressure (-Δp). Probe prefilling with octadecafluorooctane allows for optical and physical separation of the extract within the cantilever. (**B**) Selective extraction of organelle components by tuning the aperture size and thus the applicable range of applied fluidic forces. Top row: Schematic view of extracted cell components inside the cantilever. Middle row: scanning electron microscopy images of cantilever apexes with different apertures. Bottom row: Range of applicable fluidic forces with adapted FluidFM cantilevers. Scale bar. 2 µm. (**C**) Schematic of mitochondria injection into single cells by applying positive pressure (+Δp) once the cantilever was inserted into the recipient cell.

**Fig. 2.**
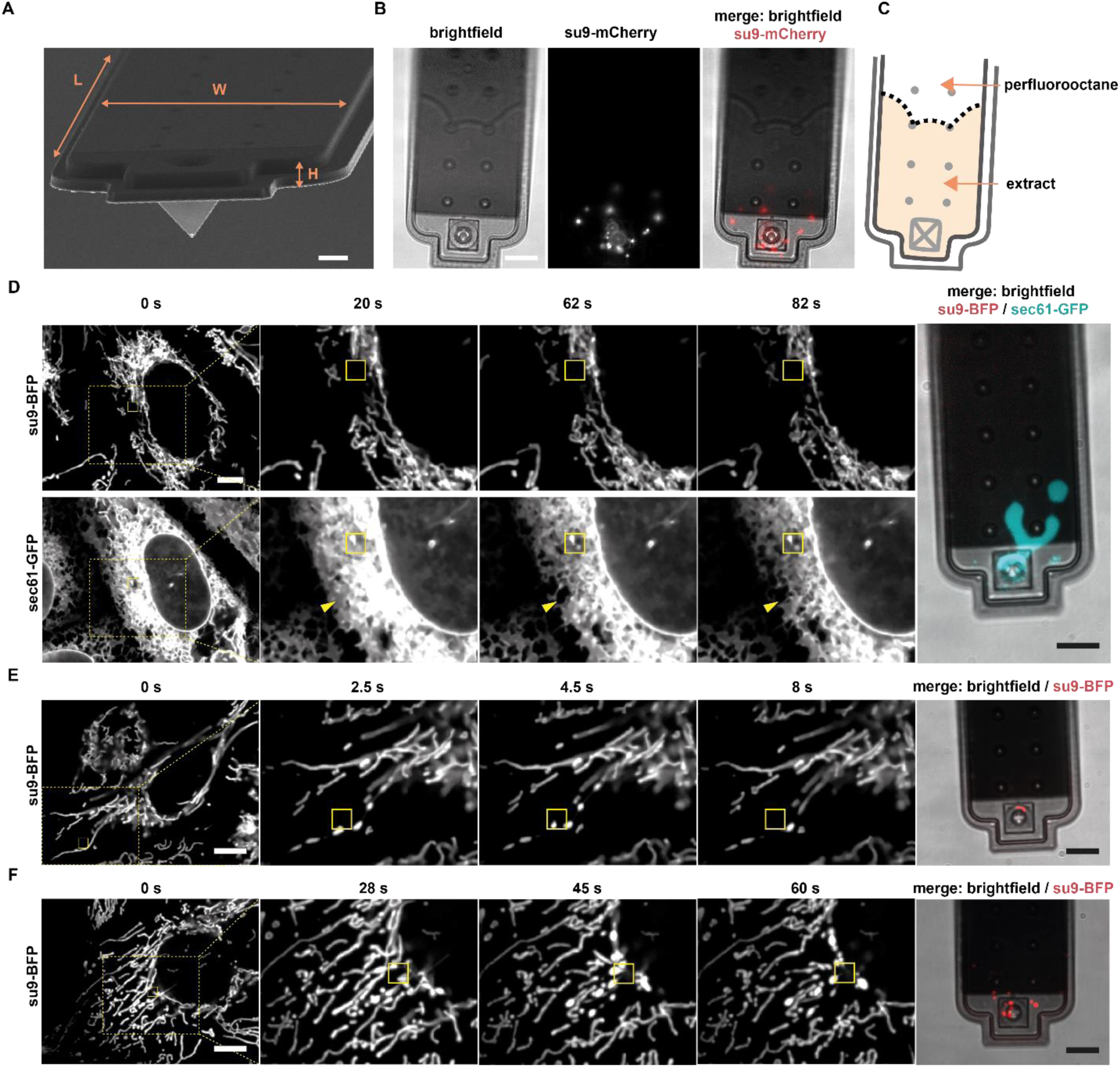
Organelle extraction. (**A**) Scanning electron microscopy image of a FluidFM cantilever. Channel dimensions: L = 200 μm, W = 35 μm, H = 1 μm. Scale bar = 5 μm. (**B**) Fluorescence microscopy images of a FluidFM cantilever after mitochondria extraction. The border between the extract and perfluorooctane can be seen due to different refractive indexes. Scale bar = 10 μm. See also: Supplementary Movie 1 (**C**) Scheme of the extract and perfluorooctane inside the cantilever shown in (B). The extract occupies a volume of 1170 fL. (**D**-**F**) Time-lapse images of live-cell compartment extractions and z-stacks of pyramidal cantilevers post-extraction. Yellow boxes indicate position of the cantilever apex inside the cell. (**D**) U2OS cell expressing su9-BFP (mitochondria) and Sec61-GFP (ER, cyan) during extraction. Arrows indicate the zone of ER retraction. A cantilever with an aperture of 0.5 µm^2^ was used. Scale bar: 10 µm. See also: Supplementary Movies 1 and 2. (**E**) Extraction of a single mitochondrion from a viable U2OS cell expressing su9-BFP. A cantilever with a 1 µm^2^ aperture was used. Scale bar: 10 µm. See also: Supplementary Movie 3. (**F**) Extraction of several mitochondria from a viable U2OS cell expressing su9-BFP. A cantilever with a 1 µm^2^ aperture was used. Scale bar: 10 µm. See also: Supplementary Movie 4.

## Results

### Tunable organelle extraction from live cells

To enable organelle manipulations, in particular their unconstrained flow through the probe, we manufactured FluidFM-probes with a channel height of 1.7 µm and drilled apertures up to 1100 nm × 1100 nm (*A* = 1.2 μm^2^) with focused ion beam (FIB) (Fig. 1B). In addition, we designed and fabricated dedicated probes with a cylindrical tip to facilitate minimally invasive cell entry. These were sharpened by FIB milling to resemble a hollow needle to facilitate membrane insertion (Fig. 1B, Supplementary Fig. 1). These probes had an aperture area of *A* = 1.6 μm^2^, further minimizing steric limitations and increasing the range of applicable hydrodynamic forces spanning from a few pN to over 100 nN, while showing great robustness towards mechanical stress (Supplementary Fig. 2). The general workflow for the manipulation of intracellular membrane enclosed compartments involves positioning the FluidFM probe above a selected subcellular location and their insertion by AFM force spectroscopy, followed by either extraction of material from the cell by exerting negative pressure (Fig. 1A, Supplementary Movie 1) or injection into the cell by positive pressure (Fig. 1C). Exclusion of large organelles is achieved by fine-tuning the aperture size (*A*) and the strength of the applied negative pressure (−*Δp*). The extraction of cytoplasmic material is monitored in real time and the extract is inspected inside the FluidFM channel by optical microscopy after relieving the pressure (*Δp* = 0) and retracting the probe (Fig. 2A-C, Supplementary Movie 1). FluidFM probes with a channel height of 1 µm can capture a total volume of 7 pL. The extraction process allows exclusion of organelle compartments that require larger apertures and higher hydrodynamic forces for extraction. However, when larger apertures are used, smaller and less strongly crosslinked organelles will be co-extracted (Fig. 1B).

The sampled material can be dispensed subsequently for downstream analyses or transplanted directly into a recipient cell (Fig. 1A). To examine the capabilities of the newly fabricated FluidFM probes for organelle sampling from single cells, we monitored the endoplasmic reticulum (ER) and mitochondria. We used human osteosarcoma epithelial (U2OS) cells and visualized in parallel the ER by expression of GFP fused to the resident protein Sec61β (sec61-GFP) and mitochondria by expression of BFP targeted to the mitochondrial matrix (su9-BFP). When utilizing pyramidal probes with an aperture size of *A* = 0.5 μ*m*^2^ and low pressure offsets, *Δp* < 20 mbar, we accumulated ER in the cantilever, which was accompanied by disappearance of GFP-signal in the cell (Fig. 2D, Supplementary Movie 2). During extraction, the ER was pulled towards the cantilever tip and we observed a general conversion of cisternal to tubular ER, in both U2OS cells and a similarly labelled kidney cell line (COS7, Supplementary Fig. 3, Supplementary Movies 2 and 3). Notably, under these conditions, the mitochondrial network remained unperturbed and mitochondria were not extracted.

Next, we aimed at extracting mitochondria using pyramidal probes with a larger aperture size (*A* = 1.2 μm^2^) and newly developed, slanted cylindrical probes (*A* = 1.6 μm^2^) (Fig. 1B). Tunable extraction of mitochondria was achieved using both kinds of probes, thus enabling aspiration of individual mitochondria or sampling of larger quantities of the mitochondrial network (Fig. 2E-F; Supplementary Movies 4 and 5).

We examined cell viability upon subcellular manipulation of ER and mitochondria and did not find it compromised (>95% cell viability) (Supplementary Fig. 4). To further ensure that our extraction protocol does not damage the cytoplasmic membrane upon probe insertion, we conducted a dedicated set of experiments and monitored potentially occurring Ca^2+^ influx from the cell culture medium using a fluorescent probe (mito-R-GECO1 (*28*)). Our experiments confirmed that there was no ion influx during and after manipulation, indicating integrity of the cytoplasmic membrane during organelle extraction (Supplementary Movies 7-8).

Monitoring mitochondrial extraction, we noticed that mitochondrial tubules exposed to tensile forces (negative pressure) underwent a shape transition reminiscent of a ‘pearls-on-a-string phenotype’ (29) inside the cytoplasm of the target cells. This phenotype was characterized by discrete spheres of mitochondrial matrix, connected by thin and elongated membrane stretches (Supplementary Fig. 5A-B). These globular structures eventually pinched off upon further exertion of a pulling force and resulted in spherical shaped mitochondria in the cantilever (Fig. 2E, Supplementary Figure 5A-C). To date, it was not possible to exert hydrodynamic forces intracellularly, while distinguishing physical manipulation from other potential cellular triggers. The observed mitochondrial ‘pearls-on-a-string’ phenotype was previously described to result from calcium overflow (29) or mitochondrial membrane rupture (30). To ensure mitochondrial membrane integrity and thus functionality, we investigated whether both mitochondrial membranes remained intact during the process. To this aim, we used U2OS cells and performed time-lapse microscopy during extraction of mitochondria in which both the matrix (su9-BFP) and the outer mitochondrial membrane (Fis1TM-mCherry) were labelled. Consistent with the observations described above (Fig. 2E), we observed the morphological change of mitochondrial tubules exclusively under direct fluid flow at the cantilever aperture and thus exertion of tensile forces, but not upon probe insertion without fluid flow (Supplementary Fig. 5A-B, Supplementary Movie 5-6). Our data further show that the force-induced shape transition propagated over tens of micrometers along the mitochondrial tubules in the millisecond to second range after negative pressure was applied with FluidFM (Supplementary Fig. 5B-C and Supplementary Movie 6). The shape transition of the matrix compartment propagated homogeneously along connected mitochondrial tubules, while the outer mitochondrial membrane (OMM) between the matrix foci initially remained intact. When traction was maintained for a few seconds, the OMM separated at one or more constriction sites between previously formed ‘pearls’, which resulted in isolated spherical mitochondria, while the remainder of the tubular structure relaxed and recovered (Supplementary Fig. 5A-C and Supplementary Movie 6). Furthermore, we demonstrate that the observed scission process of ‘pearling’ mitochondria (Supplementary Fig. 5C) succeeds recruitment of the fission machinery protein GTPase dynamin-related protein 1 (Drp1) (31) to the constricted sites (Supplementary Text and Supplementary Fig. 5D). Combining mitochondrial pulling experiments with mitochondria-localized calcium sensors, we were able to show that the shape transition towards the pearls-on-a-string phenotype and subsequent mitochondrial fission inside the cytoplasm was calcium-independent (Supplementary Fig. 6, Movies 7-10) (Supplementary text on Force-induced mitochondrial fission for details on fluorescent probes, conditions and controls, Supplementary Fig. 5 and 6, Movies 7-8).

### Mitochondrial transplantation into cultured cells

Our next goal was to demonstrate the functional delivery of mitochondria into new host cells and to achieve cell-to-cell organelle transplantation. In contrast to mitochondria extraction, for which both pyramidal probes and cylindrical probes could be used (Fig. 1B), injection of mitochondria was possible only with the latter, newly developed probes. FluidFM offers two possibilities for mitochondrial transfer: transplanting mitochondria from a donor cell to a recipient cell by coupling mitochondrial extraction with re-injection of the extract into a new host cell, or back-filling FluidFM probes with mitochondria purified by subcellular fractionation, followed by injection (Fig. 3A-D). Working with bulk-isolated mitochondria allows for a higher throughput of cells injected in series with one cantilever (> 1 cell per minute). However, such a protocol is accompanied by reduced mitochondrial quality caused by the preceding purification process. We compared both approaches, the cell-to-cell approach (Fig. 3A) and the injection of purified mitochondria (Fig. 3C), with respect to the delivery of mitochondria into the cytoplasm of individual cultured HeLa cells. To visualize the transfer of mitochondria, we used donor and acceptor cells with a differentially labelled mitochondrial matrix (Fig. 3E-F; su9-mCherry and su9-BFP respectively). When transplanting mitochondria directly from cell to cell using FluidFM, we achieved successful transfer of mitochondria into the cytosol of the recipient cells in 95% of all cases, while maintaining cell viability (Fig. 3G, 39 out of 41 transplanted cells). Upon injection of purified mitochondria, we observed mitochondrial transfer and preserved cell viability in 46% of cases (19 of 41) (Fig. 3G, Supplementary Fig. 7). Quantification of the transplant showed that the number of transplanted mitochondria for these experiments varied from 3 to 15 mitochondria per cell (Fig. 3H). The different success rates between the two alternative protocols can be explained by differences in mitochondrial condition. When evaluating mitochondrial extraction protocols, we observed that a fraction of the extracted mitochondria undergo rupture of their outer membranes (Supplementary Fig 8) (32). Irreversible damage of mitochondria leads to degradation inside cells and potentially cell apoptosis due to cytochrome c leakage. While cell-to-cell transplantation of mitochondria reduces throughput, it has the advantage that the extracellular time is short (< 1 min) and that mitochondria sampled by FluidFM are maximally concentrated in native cytoplasmic fluid, bypassing the use of artificial buffers altogether. We ensured that the extract remained near the aperture during extraction by filling the probes with immiscible perfluorooctane before extraction and transplantation. Therefore, only small volumes (0.5 – 2 pL) are injected into the host cells (Fig. 3B), up to the volume previously extracted from the donor cell (injection of larger volumes is automatically prevented due to inherent flow resistance properties of the pre-filled fluorocarbon liquid).

**Fig. 3.**
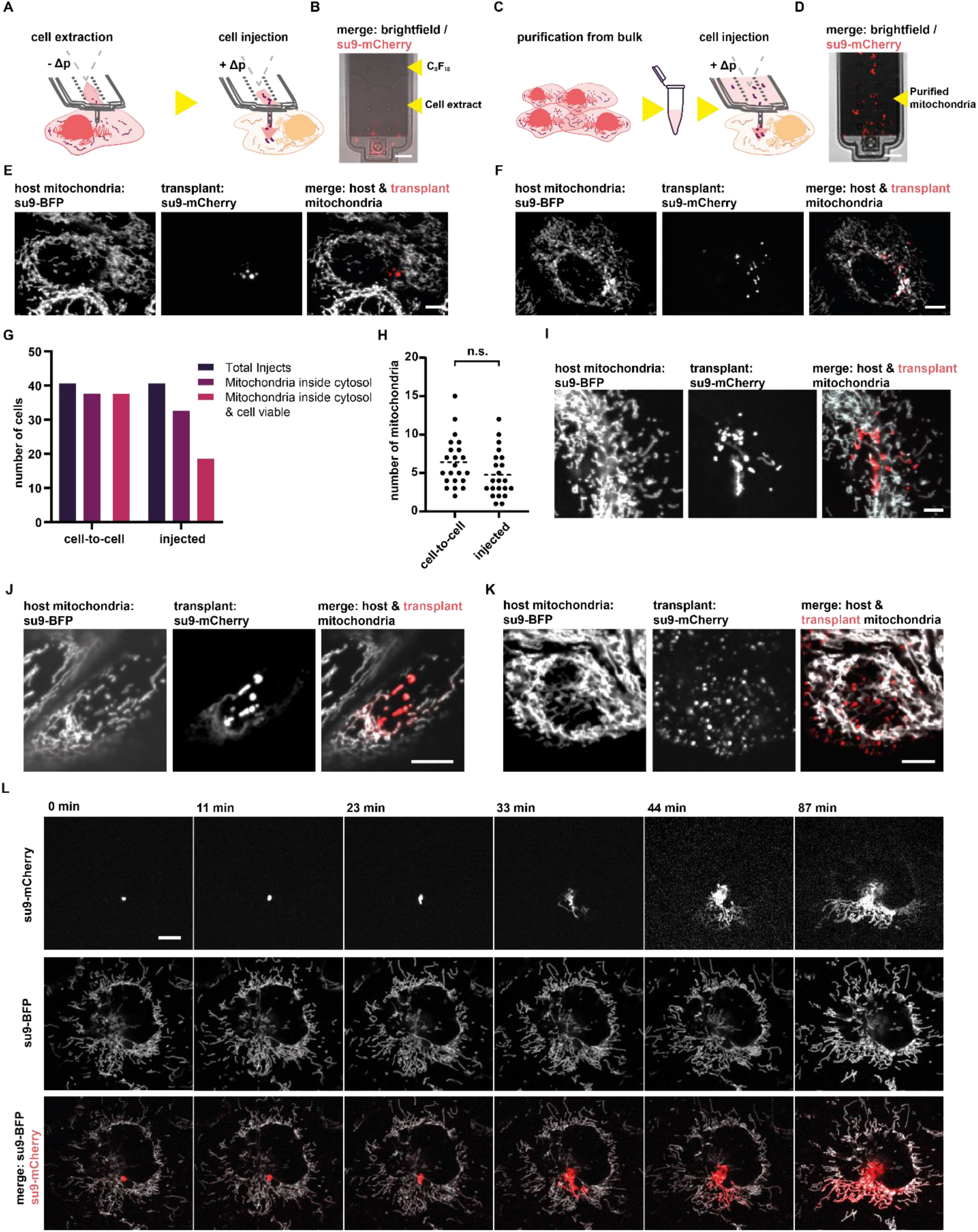
Mitochondrial transplantation. (**A**) Scheme of mitochondrial transplantation using the cell-to-cell transfer approach: Mitochondria are extracted via FluidFM-aspiration. Subsequently, the cantilever holding the extract is moved to a recipient cell and the extract is injected. (**B**) Image of a FluidFM cantilever prefilled with perfluorooctane after mitochondrial extraction, mitochondria are labelled via su9-mCherry. Extracted volume ∼0.8 pL. Scale bar: 10 µm. (**C**) Scheme of mitochondrial transplantation using mitochondria purified by a standard mitochondrial purification protocol. The purified mitochondria are resuspended in HEPES-2 buffer and filled directly into the fluidic probe. Cells are injected consecutively. (**D**) Image of a FluidFM cantilever filled with mitochondria isolated from bulk, labelled via su9-mCherry. Scale bar: 10 µm. (**E**) Images of a recipient cell post mitochondrial transplantation via the cell-to-cell approach. The host cells’ mitochondrial network is labelled via su9-BFP and the transplant is labelled via su9-mCherry. Scale bar: 10 µm. (**F**) Images of a recipient cell post mitochondrial transplantation via the injection of isolated mitochondria approach, labels similar to c. Scale bar: 10 µm. (**G**) Evaluation of mitochondrial transplantation via the cell-to-cell approach upon optical inspection and the injection of isolated mitochondria approach. 40 cells were evaluated per approach. (**H**) Absolute numbers of transplanted mitochondria of 22 individual cells evaluated for the cell-to-cell and the injection of isolated mitochondria approach. (**I**) Fusion states of transplanted mitochondria 30 post cell-to-cell transplantation. Mitochondria are visualized using different fluorescent labels for the transplant (su9-mCherry) and for the host mitochondrial network (su9-BFP). Scale bar: 5 µm. (**J**) Fusion states of transplanted mitochondria 30 post injection of purified mitochondria, similar labelling as in g. Scale bar: 5 µm. (**K**) Degradation of transplanted mitochondria, the transplant is split into multiple smaller fluorescent vesicles (su9-mCherry) showing no overlap of fluorescence with the labelled host cell mitochondrial network (su9-BFP). Scale bar: 5 µm. (**L**) Time lapse image series of a single transplanted mitochondrion (su9-mCherry). The organelle donor was a HeLa cell, recipient cell is a U2OS-cell with a fluorescently labelled mitochondrial network (su9-BFP). Scale bar: 10 µm.

Labeling mitochondria of the recipient cell (su9-BFP) in addition to labeling donor cell mitochondria (su9-mCherry) allowed us to survey the state of the mitochondrial network in the transplanted cell. In both transfer approaches described above (transplantation and purification followed by injection), the tubular, interconnected phenotype of the host-mitochondrial network remained unaltered by the injection process. In addition, labeling allowed us to monitor the fate of the transplanted mitochondria. We observed mitochondrial acceptance, defined by fusion of the transplant with the host mitochondria network, and thus overlap of both fluorescent signals- and mitochondrial degradation, marked by further fragmentation of the transplant and segregation into presumptive mitophagosomal structures (Fig. 3I-K). These processes were observed irrespective of the transfer method, cell-to-cell (Fig. 3I), or injection of purified mitochondria (Fig. 3J). We followed the fate of the transplant over time in 22 cells: 18 cells showed full mitochondrial fusion of the transplant and 4 cells showed mitochondrial degradation. Fusion events were first observed within 30 minutes post transplantation in the majority of cases (14 of 18 cells).

As indicated above, the high cell-to-cell transplant efficiency allows for direct observation of the fate of individually transplanted mitochondria. To showcase this, we transplanted labelled mitochondria (su9-mCherry) from HeLa cells into differentially labelled U2OS cells (su9-BFP), regularly used for studies of dynamic mitochondrial behavior. The strong label together with a highly sensitive EMCCD-camera (Andor) enables tracing of individual mitochondria within a recipient cell over time (Fig. 3I, Supplementary Movie 11). In this particular case, we observed rapid spread of the fluorescent mitochondrial matrix label after the initial fusion event to the network 23 minutes post transplantation.

In summary, we established two methods for mitochondria transfer into single cultured cells. One involves bulk purification of mitochondria and their injection into recipient cells. The injection protocol is rather rapid but inevitably compromises mitochondrial and cellular function. The second consists in cell-to-cell transplantation. Evaluation of cell viability and transplant fate showed an efficient protocol that allows observation of the dynamic behavior of transplanted mitochondria after transfer.

### Fate of transplanted mitochondria in primary cells

Having developed an efficient protocol for cell-to-cell transplantation of mitochondria, we sought to test whether primary cells show similar uptake behavior as the tested cancer cells and, if so, what are the dynamics of integration of foreign mitochondria. We considered these particular experiments important because quality control mechanisms are impaired in cancer cell lines (33) and to demonstrate the broad applicability of the established protocol. In addition, several studies link naturally occurring mitochondrial transfer events with short-term benefits for individual cells and tissues, for example in osteocytes (34), adipose tissue (35) or in neurons (36). However, to the best of our knowledge, the fate of mitochondria or dose-response relationships have not be studied, and appropriate technologies of mitochondrial transfer that preserve cell viability have been lacking.

We used primary human endothelial keratinocytes (HEKa), a skin cell type that is generally susceptible to radiation damage and aging (37). In standard culture conditions, the mitochondrial network of HEKa cells is mostly tubular, forming a large connected network (Supplementary Fig. 8) similar to HeLa cells studied above, indicating an active mitochondrial fusion machinery (38). Notably, cell-to-cell transplantation of mitochondria into HEKa cells showed no impact on their viability, allowing analysis of all injected cells. We conducted time-lapse experiments of transplanted labelled mitochondria (su9-mCherry) from HeLa cells into HEKa cells, focusing on the fluorescent signal of the transplant and its dynamic behavior over time (Fig. 4A-C Supplementary Movie 12). The analysis revealed the rate of transplant fusion with the host mitochondrial network and its movement inside the cytosol. In contrast to the findings described above for HeLa-to-HeLa transplantations, i.e. host cells either accepting or degrading mitochondria (Supplementary Fig. 9), we observed a third, intermediate scenario in which single cells both fused-and degraded parts of the transplant. To further define the transplant uptake mechanisms in HEKa cells, we classified the transplanted mitochondria as primary particles – mitochondria that retain their original shape and fluorescence intensity, indicating that they have neither fused with the host network, nor been targeted to degradation –, secondary particles – mitochondria that retain their fluorescence but are distinctively smaller in size, suggesting fragmentation for degradation –, and tertiary particles – mitochondria fused with the host network, showing a characteristic decrease of transplant specific fluorescence (> 10 fold), due to dilution into the host mitochondrial network (Fig. 4D, E, Supplementary Movie 12). Shape transition of the transplant from discrete spheres towards tubules, was observed exclusively in the context of fusion of the transplant to the existing host network. Emergence of secondary particles hinted towards rearrangement of the transplant into parts recognized as ‘viable’ mitochondria, designated for fusion to the network, while another part was targeted for degradation. Tracing this process within an individual cell, we counted the number of ‘primary’ and ‘secondary’ particles over 14 h (Fig. 4F). In the first 3 h post injection, we observed an increase in ‘primary’- and ‘secondary particles’, which subsequently lost their high fluorescent signature while the fraction of fluorescent mitochondrial network increased over time (Fig. 4B-E). To test whether restructuring of the transplant into sub-particles was common in cells that showed emergence of secondary particles, we counted the number of primary and secondary particles immediately after transplantation- and 3 h after in cells that showed both degradation and integration of the transplant (Fig. 4G; n = 10). The results indicated variance in the absolute extent of rearrangement; however, all cells showed an increased particle number by a factor of 1.7 ± 0.4.

**Fig. 4.**
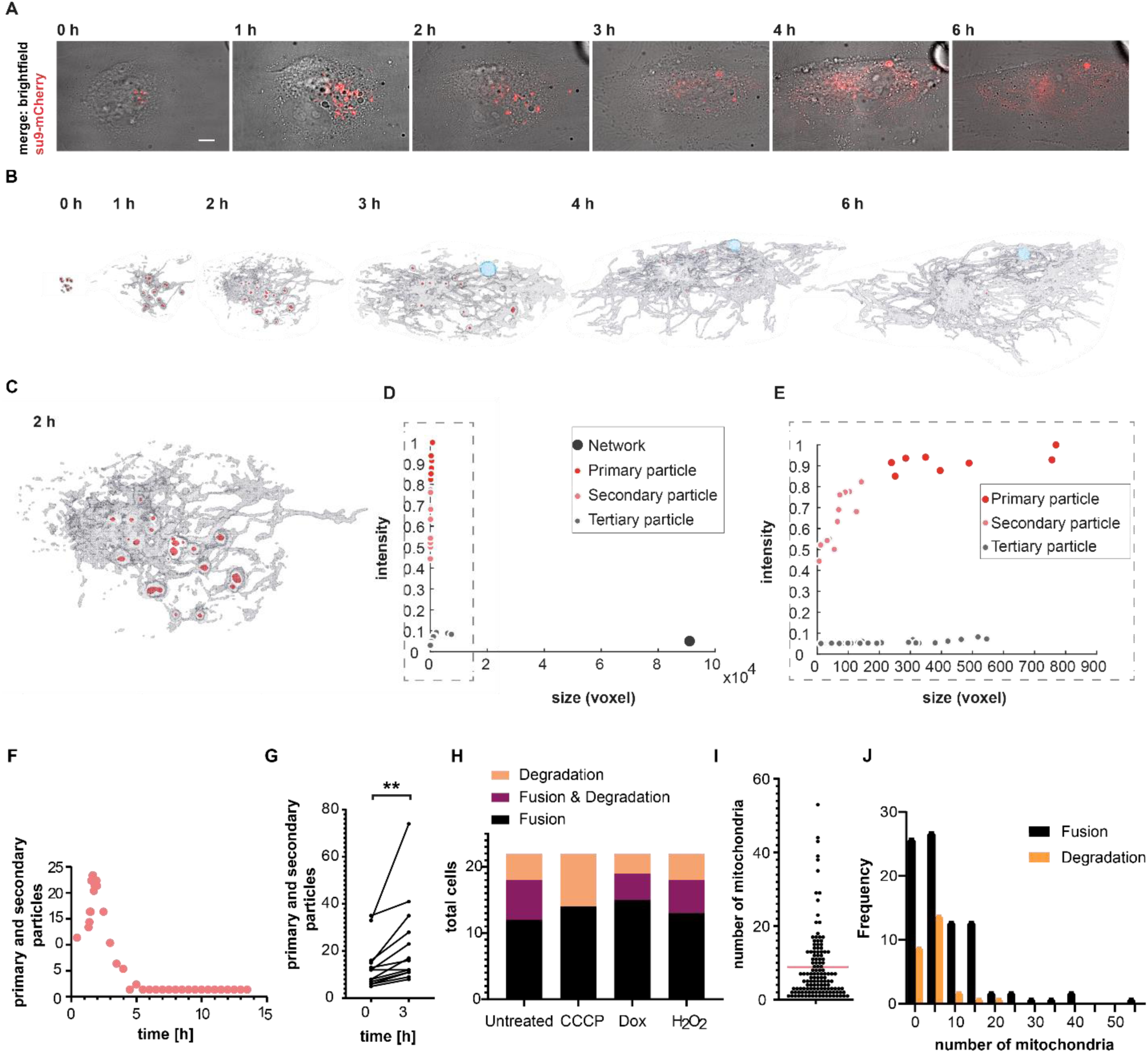
Mitochondrial fate in HEKa cells. (**A**) Time series of cell-to-cell transplantation from HeLa (su9-mCherry) to HEKa, fusion and degradation to the unlabeled host cell network. Scale bar: 10 µm. (**B**) Surface-render (total su9-mCherry signal) of the time-lapse series of mitochondrial fusion and degradation within the host cell shown in a. The transplant is depicted in red, host cell mitochondrial network carrying fluorescent traces of the transplant is shown in grey, presumptive mitophagosomal structure is shown in light blue. (**C**) Enlarged section of the 2 h timepoint surface render from b. (**D**) Scatter plot of size and normalized fluorescence intensity distribution of objects detected at the 2 h-timepoint. (**E**) Zoom in on low-size fraction of the scatter plot shown in c. (**F**) Total number of objects classified as primary transplant or secondary particle over time. (**G**) Absolute number of primary and secondary particles detected directly after transplantation (0 h) and after 3 hours for 12 cells. Paired t-test, p = 0.0095. (**H**) Fusion and degradation behavior of drug-compromised mitochondrial transplants in HEKa cells. Each condition was tested with 22 cells. (**I**) Distribution of mitochondrial quantity transplanted per cell from all conditions tested in e, total number of cells: 132. Dashed line indicates mean value. Total number of transplanted mitochondria: 1117. (**J**) Histogram plot correlating the amount of transplanted mitochondria with uptake outcome; bin width is 5. Correlation of the absolute number of transplanted mitochondria per cell with the cell response of either mitochondrial fusion- or degradation across all transplant conditions in percent. Total cell number: n = 135.

In this first set of transplantation experiments from HeLa cells to HEKa cells, 12 cells showed complete uptake of the transplant, 6 showed both uptake and degradation of the transplant, and 4 fully degraded the transplant (Fig. 4H). The degradation of defined amounts of transplanted mitochondria can be traced over time in single cells, as exemplarily shown in Supplemental Figure 10.

In the experiments described above, the first fusion or degradation events occurred 20 minutes post transplantation and continued for more than 16 hours. Remarkably, the speed at which the primary transplant was processed was independent of transplant sizes. Fusion or degradation of large transplants, >40 mitochondria per cell, advanced at similar rate as smaller transplants, <8 mitochondria per cell (Supplementary Movie 12, Supplementary Fig. 11).

As outlined above, the established cell-to-cell transplantation protocol is minimally invasive with regard to the integrity of the mitochondria themselves when using ‘healthy’ donor cells, and cells receiving transplants from unperturbed donor cells showed mostly uptake of the transplant (Fig. 4H). Next, we wondered whether the acceptance of mitochondria by host cells was altered, if the quality of the transplant was impaired by prior drug treatment of the donor cell. Such treatments are commonly used to study the pathways that control the maintenance of the mitochondrial network. However, drugs have been applied to the entire cell so far, likely interfering with regulatory processes of cells as a whole. In consequence, the here established cell-to-cell transplantation of mitochondria provided an opportunity to follow the fate of damaged mitochondrial sub-populations in the context of an otherwise intact cell. *In vitro*, the chemical triggers used to study mitophagy are uncoupling agents, inhibitors of the respiratory chain or combinations thereof, robustly causing changes of membrane potential and simulating low mitochondrial quality (39). To investigate how otherwise healthy cells respond to impaired mitochondrial subpopulations, we treated transplant-donor cells with the proton ionophore CCCP (Supplementary Fig 8) and transplanted the depolarized mitochondria into HEKa cells. The majority of cells, 14 of 22, showed fusion of initially depolarized mitochondria to the network, while 8 cells degraded the transplant (Fig. 4H). This reaction was similar to the previous condition tested, the transplantation of untreated mitochondria, which was unexpected, because literature suggests rapid mitochondrial degradation of mitochondria having lost their membrane potential (40, 41). However, the membrane potential can potentially recover quickly, if a functional ATPase is present, as expected in our study. Next, we tested the influence of Doxycycline, an antibiotic, inhibiting protein synthesis of the mitochondrial ribosome, which induces fractionation of the mitochondrial network in HeLa cells after 24 h (Supplementary Fig. 8), putatively due to non-functional mitochondrial OXPHOS protein complexes (42). The reaction of the host cells was similar as towards depolarized and untreated mitochondria: 15 show full fusion, 4 show both fusion and degradation and 3 show full degradation of the transplant. We then treated donor cells with H_2_O_2_, which damages proteins and induces double strand breaks in mtDNA. The fraction of mtDNA containing double strand breaks in H_2_O_2_ treated cells was reported to remain high for several hours post treatment (43) and the donor cells showed a fragmented mitochondrial network after 3 h (Supplementary Fig. 8). However, even upon H_2_O_2_ treatment, and even upon application of all drugs simultaneously to donor cells, mitochondrial acceptance in healthy recipient background was still comparable to all other conditions, indicating the potential of cells to cope with highly damaged mitochondria when occurring as isolated events (Fig. 4H, Supplementary Fig 12).

Since the tested conditions for drug-impaired mitochondria did not show an impact on mitochondrial uptake behavior, we wondered whether the amount of injected mitochondria might have an impact on transplantation outcome. Across all conditions, we transplanted 1117 mitochondria (Fig. 4I) and followed their fate in more than 100 individual primary cells after transplantation of 1 up to 53 mitochondria (Supplementary Fig. 12). Pooling uptake data of cross-tested conditions (n = 135 cells), we plotted the frequency of cells showing transplant fusion and transplant degradation vs the number of individually transplanted mitochondria and could not find any correlation between the number of transferred mitochondria per cell and their uptake behavior (Fig. 4J).

Overall, transplantations were successful in all experiments and cells remained viable, irrespective of the amount of transplant received. We demonstrated that primary HEKa cells incorporate a majority of transplanted mitochondria into their network via mitochondrial fusion. A subset of transplanted cells showed individualized responses, with one fraction of mitochondria fusing with the mitochondrial network and another subpopulation undergoing rearrangement associated with particle formation, while a third fraction was subjected to degradation. This behavior highlights the presence of quality control mechanisms that are in place to sort each mitochondrion and exemplifies the potential of the FluidFM approach to transplant mitochondria into new host cells to study their fate and host cell response with single mitochondria resolution.

### Mitochondria transplantation and transfer of mitochondrial genomes

After demonstrating the short-term response of cells to mitochondrial transplantation, we focused on the long-term effects of mitochondrial transfer over generations of host cells. Mitochondria differ from other membrane compartments in that they carry their own genome, which is propagated within the cell’s inherent mitochondrial pool. It has been shown that mtDNA can be transferred into somatic cells via miniaturized probes under selective pressure, either by transfer into cells artificially rendered free of mitochondrial DNA (rho-zero cells) (22, 44) (2-25% efficiency), or by selection using antibiotics (< 0.01% efficiency) (10). Therefore, the introduction of new mtDNA sequences into functional somatic cells remains a challenge (45, 46). Before demonstrating the transfer of mtDNA into host cells upon transplantation, we first wondered whether the FluidFM-extracted mitochondria contain mitochondrial DNA, which is organized in discrete complexes termed mitochondrial nucleoids, because the extraction process leads to rapid fragmentation of the network. To visualize the behavior of mitochondrial nucleoids during mitochondrial extraction using FluidFM, we expressed a fluorescently tagged version of p55 (p55-GFP), a polymerase-γ subunit that co-localizes with mitochondrial nucleoids and appears in discrete speckles scattered throughout the mitochondrial matrix (47) (Fig. 5A). We evaluated time-lapse experiments upon mitochondrial extraction with labelled nucleoids and counted mitochondrial fragments either containing labelled nucleoids, or being devoid of their fluorescent signal (Fig. 5B and Supplementary Fig. 13). We observed p55-GFP in > 90% of formed mitochondrial spheres (n = 18), indicating that most transplanted mitochondria contained mtDNA.

**Fig. 5.**
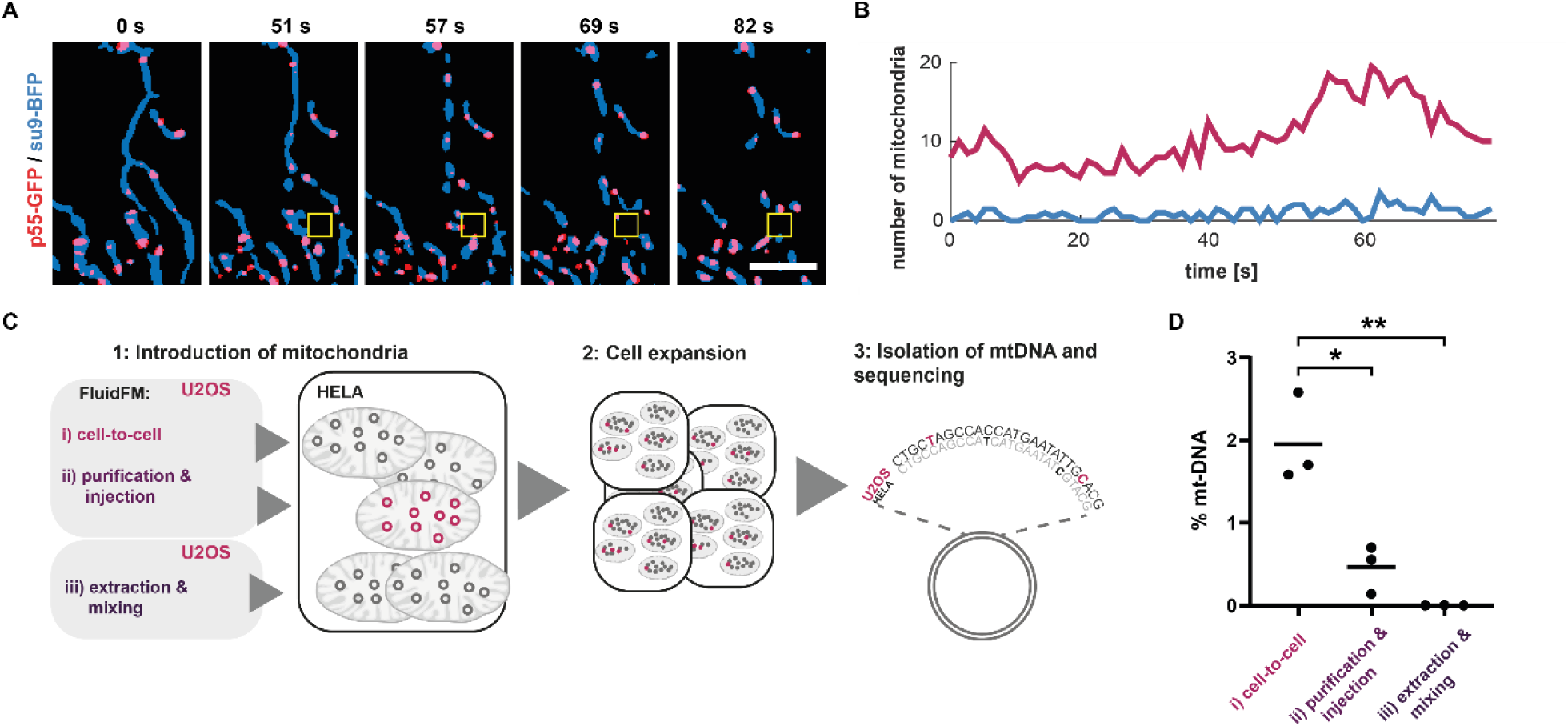
Mitochondria transplantation and transfer of mitochondrial genomes. (**A**) Time-lapse evaluation of mitochondria extraction visualizing mitochondrial nucleoids. An overlay of mitochondrial matrix shown in blue (su9-BFP) and mitochondrial nucleoids shown in red (p55-GFP). Yellow box indicates position of the cantilever. **(B**) Dynamic localization of labelled nucleoids within a fractionating mitochondrial network. Plotted are the total amounts of mitochondria (su9-BFP) that show overlap with a labelled nucleoid (p55-GFP) in red and idle mitochondria showing no fluorescent trace of p55-GFP in blue over time. See also Supplementary Fig. 13. Scale bar: 5 μm. (**C**) Strategy for the quantification of mtDNA maintenance after mitochondrial transplantation. (**D**) Quantification of retained U2OS mtDNA in transplanted HeLa cells of all approaches tested. Bars indicate mean value. Difference between condition i and ii, p = 0.0138 and between condition i and iii, p = 0.0034.

Next, we applied genomics to quantify uptake and maintenance of mtDNA after mitochondria transplantation (Fig. 5C). We compared two methods of mitochondrial transfer, using U2OS cells as mitochondria donors and HeLa cells as acceptors: direct cell-to-cell transplantation of mitochondria with FluidFM and FluidFM-injection of purified mitochondria. In addition, to control for non-specific uptake of mitochondria or otherwise unspecific mtDNA carryover, we seeded cells into a culture-dish and mixed them with extracted mitochondria from donor cells. All experiments were executed in biological triplicates and cells were passaged onto a fresh culture dish, for a more detailed description of the experimental procedure and conditions, see Supplementary information. Subsequently, we extracted DNA, then amplified and sequenced part of the variable D-loop region of the mitochondrial genome. Four single nucleotide polymorphisms (48) allowed the detection of transplanted mtDNA. The percentage of transplant mtDNA propagation was highest in samples from cell-to-cell mitochondria transplantation (mean 2 %), followed by FluidFM injection of purified mitochondria (mean 0.5%), whereas mtDNA transfer by non-specific uptake was below the detection limit (Fig. 5D and Supplementary Table 1 & 2). The obtained amount of mtDNA was well in line with our expectations considering the amount of mtDNA copies per cell (∼1100) (49) and mtDNA copy number per nucleoid (1.4-3.2) (50, 51), and the quantity of mitochondria we transplanted within this experimental series (Fig. 3H, median = 6).

In summary, we showed efficient transplantation of mitochondrial DNA into somatic cells without the need for selective pressure. These results indicate that HeLa cells show little or no selection for mitochondrial DNA variants of another cell line, here U2OS cells, under standard culture conditions.

## Discussion

The technology developed here allows the manipulation of intracellular compartments using FluidFM. We show the removal and injection of organelles from and into single cells without compromising organelle integrity nor cell viability. The scalability of the volume of the subcellular sampling protocol for organelles allows molecular downstream analyses, which are increasingly feasible thanks to the improving sensitivity of ‘omics’ technologies (52). Samplings can be performed at one time point, but also repeatedly from one cell to unravel the dynamic behavior within individual cells. We demonstrate the application of FluidFM to exert localized fluidic forces within single cells and thus expand the possibilities for subcellular sampling as well as the study of organelle mechanobiology (17, 26). It has been proposed that mechanical forces and membrane constriction affect mitochondrial shape and dynamics (53, 54), but with the previously established tools it has been difficult to test such a hypothesis without perturbing the cellular state as a whole (30, 53). Gonzalez-Rodriguez et al (30) suggest that mitochondrial shape depends on the elastocapillary number (*Nec*) (55) describing the ratio between the elastic modulus (*E*) and mitochondrial membrane tension (*γ*): *Nec* ≡ *E* × *d*/*γ* where d is the membrane thickness. A decreasing *Nec* results in ‘pearling’ (30, 55). This description explains our observations where pulling on the mitochondrial membrane (Supplementary Fig. 5A) increases the mitochondrial membrane tension, thus decreasing *Nec*. While the application of compressive force can be controlled in time and space (56), application of controlled tensile force has been impossible to date. Here, we demonstrate that mechanical force can be a driver for mitochondrial shape transition that is strictly localized to sites of tensile force application and propagates along membrane-connected mitochondrial tubes. Here, the purely mechanical nature of FluidFM presents itself as a strength, because it allows dissection from complex physiological stimuli, often involving calcium-influx, from isolated exertion of (hydrodynamic) pulling forces. ‘Pearling’ eventually leads to recruitment of Drp1 and mitochondrial fission. In a more physiological setting, the transition into the pearls-on-a-string phenotype appears to be an elegant solution that protects cells against membrane leakage during mechanical stress.

Transplantation of mitochondria at a high efficiency allowed us to track organelle fate over time in new genetic and physiological cell backgrounds. Similar to organ transplants that are accepted or rejected by new hosts, here we show rescue or failure of organelles within single cells after transplantation. We show that transplantation is highly efficient (95% success rate) when mitochondria are transplanted directly between living cells rather than using isolation protocols prior to transfer. The FluidFM-based approach of efficient mitochondria transplantation permitted us to evaluate mitochondrial quality control in primary cells by transplanting healthy and compromised mitochondria and observing their fate. We show that individual cells generally differentiate between individual mitochondria, but do not display apparent responses to mitochondria previously exposed to the tested stresses. Across conditions, we observe that the majority of mitochondria become integrated into the host network and that transplanted mitochondria give rise to secondary mitochondrial particles. Such particles are reminiscent of mitochondrial-derived vesicles that take part in mitochondrial quality control (57, 58). This indicates that the extent of mitochondrial quality control may depend on the general cellular state rather than the actual quality of the mitochondrial network. The study of mitochondrial quality control is of great interest, and the approach introduced here has the potential to significantly contribute further to this field by allowing defective mitochondria to be introduced locally in an otherwise functional cellular background. In addition, it will be interesting to study the impact of mitochondrial transplantation to the host cells considering metabolic activity and signaling responses. To date, it is not clear whether mitochondria transferred *in vivo* function as ‘exchangeable cellular factories’ that benefit host cell metabolism as donors of high-quality mitochondrial DNA, as cellular messengers or possibly in combination of the named effects. The manipulation and observation of mitochondrial subpopulations within a cell is important to drive discoveries in the dynamics e.g. of asymmetric cell division (59, 60). We show here that subcellular manipulation using FluidFM extends the scope of such studies beyond the application of optogenetic tools and enables the transplantation and observation of tunable quantities of mitochondrial subpopulations in single cells.

We tracked the short term-fate of mitochondrial subpopulations using fluorescent microscopy, but our protocol also allows for the study of the long-term impacts of mitochondrial transfer by showing the introduction of novel mtDNA variants into cell populations. While mitochondrial transfer into oocytes has been demonstrated (61, 62), there are only few mechanistic insights into mtDNA selection in eukaryotic cells, with recent notable exceptions (63, 64). Heteroplasmic cells were generated by injection of isolated mitochondria into large oocytes, reaching levels of 7% (64) of transplanted mtDNA under nonselective conditions, which is comparable with our method for somatic cells (up to 2.5%). With a spread from one up to about 50 transplanted mitochondria per cell and the opportunity to inject individual cells repeatedly, higher shares of heteroplasmic variants could be achieved. FluidFM thus represents a promising way to decipher modes of mtDNA selection in cell culture models. It will enable the identification of metabolic and genetic factors that impact shifts in mtDNA heteroplasmy and nuclear-mitochondrial crosstalk. Recently, mtDNA polymorphisms have been used to track cells in vivo (65). The FluidFM approach can directly introduce such polymorphisms, providing a genetic marker to track mtDNA within cells in complex settings.

In the future, the technique introduced here will stimulate applications in additional research areas, for example, the rejuvenation of cells with low metabolic activity in stem cell therapies (4, 5) or as an alternative strategy in mitochondrial replacement therapy approaches. Beyond, it offers new perspectives to address fundamental questions in cell biology, mechanobiology and cell engineering.

## Materials and Methods

### Fabrication process of FluidFM probes with a cylindrical tip

The process starts with a standard 4” silicon wafer in the (100) crystallographic orientation. A 400 nm thick silicon-rich nitride (SiRN) layer is deposited by Low Pressure Chemical Vapor Deposition (LPCVD) on the selected wafer. The thickness of the SiRN layer determines the thickness of the bottom wall of the microfluidic channel. A silicon oxide layer is deposited by LPCVD from TEOS on the top of the SiRN layer. Then, a circular opening is patterned by Reactive Ion Etching (RIE) on the deposited multilayer. By using Deep Reactive Ion Etching (DRIE), a cylindrical pattern is subsequently formed in silicon. The depth of the cylindrical mold defines the height of the cylindrical tip. After formation of the cylindrical mold, a 150 nm thick SiRN layer is deposited by LPCVD. The thickness of the second SiRN layer determines the thickness of the walls in the cylindrical tip. After the tip molding, a blanket etch (RIE without an etching mask) RIE is performed to remove silicon nitride from the top surface of the wafer and from the bottom of the cylindrical mold. Due to the directionality of the RIE process, the material on the side-walls is preserved. The silicon oxide layer, which protects the underlying SIRN layer during the RIE etching, is removed. Subsequently, a 1500 nm thick polysilicon layer is deposited by LOCVD. Polysilicon is used as a sacrificial material to form a microfluidic channel. The thickness of the polysilicon layer determines the height of the microfluidic channel. The layout of the microchannel is patterned by RIE of polysilicon. A SiRN layer with a thickness of 400 nm is deposited by LPCVD after the patterning of polysilicon. This SiRN layer forms the top wall of the fluidic microchannel. The layout of the cantilever and the inlet of the microchannel are defined by RIE of silicon nitride. Next, a reflective metal layer is deposited and patterned on the cantilever. The silicon wafer is anodically bonded to a pre-diced glass wafer in which the fluidic inlets are pre-manufactured by powder blasting. After the anodic bonding, the sacrificial polysilicon layer is removed by TMAH etching in order to empty the microchannel and form a hollow cantilever. The bulk silicon is also removed during the TMAH etching to completely release the cantilever. A small part of silicon remains underneath the microfluidic inlet to provide mechanical strength for the channel walls.

### FluidFM probe processing and FIB-SEM imaging and milling

The FluidFM-probes were mounted into a custom probe holder and coated with an 18 nm carbon layer using a CCU-010 Carbon Coater (Safematic) before milling by a FIB-SEM Nvision 40 device (Zeiss) using the Atlas software (Zeiss).

Pyramidal probes used for extraction of organelles and hydrodynamic pulling experiments: The milled face of the pyramidal probe was aligned perpendicularly to the FIB-beam equipped with a gallium ion source. Subsequently, the probes were milled with an acceleration voltage of 30 kV at 10 pA until the aperture was milled, controlled by optical observation (approximately to an electric charge of 10 nC per µm^2^). The pyramidal probes have a 100 nm thick silicon nitride layer at the aperture region.

Cylindrical probes used for mitochondrial transplantation experiments were ‘sharpened’ by milling the probe apex at a 50° angle alongside the cylinder at an acceleration voltage of 30 kV at 40 pA. The probes were optically controlled after the milling procedure.

Then, the probes were glued onto a cytoclip holder by Cytosurge. Before each experiment, the cantilevers were cleaned by a 90 s plasma treatment (Plasma Cleaner PDG-32G, Harrick Plasma) before coating overnight with vapor phase SL2 Sigmacote (Sigma-Aldrich) in a vacuum desiccator. The siliconized probe was oven dried at 100 °C for 1 h. The cantilever spring constant was measured using software-implemented scripts (cylindrical probes: 2 ± 0.4 Nm^-1^, pyramidal probes: 5 ± 1 Nm^-1^).

### FluidFM setup and Microscopy

The FluidFM setup is composed of a FlexAFM 5-NIR scan head controlled by a C3000 controller (Nanosurf), a digital pressure controller (ranging from -800 mbar to +1000 mbar), and Microfluidic Probes (Cytosurge). The scan head is mounted on an inverted AxioObserver microscope equipped with a temperature-controlled incubation chamber (Zeiss). The microscope is coupled to a spinning disc confocal microscope (Visitron) with a Yokogawa CSU-W1 scan head and an EMCCD camera system (Andor). For all images and videos, a 63× oil objective with 1.4 numerical aperture and a 2× lens switcher was used (without lens switcher: 4.85 pixel/micron and 9.69 pixel per micron with lens switcher); images are in 16 bit format. Image acquisition was controlled using the VisiView software (Visitron); linear adjustments and video editing were made with Fiji(*66*); additionally, images and videos were noise-filtered using the wiener noise filtering function (wiener2; 3 by 3 neighborhood size) in MatlabR2018a (MathWorks). Movies were created using a self-written Matlab script in order to visualize several sections or channels within the same movie. Colormaps originate from Thyng et al.(67). Images of cantilevers containing extracts (Fig. 2) were created summing up the slices of a Z-stack via Fiji and reconverting the image to 16 bit format.

### Cell Culture

U2OS, COS7, and HeLa cells were maintained in Dulbecco’s Modified Eagle Medium containing 1% penicillin-streptomycin (ThermoFisher) and 10% fetal bovine serum (ThermoFisher) culture medium at 37°C and 5% CO_2_ in a humidified incubator. Primary adult human endothelial keratinocytes (HEKa) cells were purchased from ThermoFisher. HEKa cells were cultivated in EpiLife™ Medium, with 60 µM calcium, with Human Keratinocyte Growth Supplement, 1%, (ThermoFisher) and Gentamicin/Amphotericin Solution (ThermoFisher). Cells were seeded 48 h preceding the experiments onto 50-mm tissue-culture treated low µ-dishes (ibidi) inside two-well culture inserts (ibidi) or, for experiments coupling mitochondrial extraction via FluidFM to injection of mitochondria, inside 4-well micro-inserts (ibidi). For the experiments, the culture media was replaced with CO_2_-independent growth medium containing 10% FBS (ThermoFisher) and 1% penicillin-streptomycin (ThermoFisher). Cell lines stably expressing fluorescent proteins markers were created via lentiviral transduction, the constructs were previously described in Helle et al. (53).

### Transient transfection

U2OS-cells were seeded 72 h preceding the experiment into two-well culture inserts (ibidi) inside 50-mm tissue-culture treated low µ-dishes (ibidi) in a total volume of 100 µL per well. Transfections were performed overnight 36 h preceding the experiment using 0.2 µg plasmid DNA and 0.2 µL Lipofectamine P3000 solution following the manufacturer’s instructions (ThermoFisher). The media was exchanged to standard culture medium 12 h post transfection. The plasmid used for calcium imaging CMV-mito-R-GECO1 was a gift from Robert Campbell (Addgene plasmid #46021; http://n2t.net/addgene: 46021; RRID: Addgene_46021).

### Mitochondrial pulling and extraction experiments

All experiments were executed at 37°C. The cells for extraction/transplantation were selected by light microscopy. Z-stacks were taken before and after the manipulation step to document the workflow. FluidFM probes were prepared as specified above and prefilled with octadecafluorooctane (perfluorooctane) (Sigma-Aldrich). Subsequently, the FluidFM probe was moved over a targeted area in the cytosol of a selected cell, usually close to the nucleus or mitochondrial tubes in the cell periphery. The cantilever was then inserted at the specified location driven by a forward force spectroscopy routine in contact mode, until the setpoint of 400 nanoNewtons (nN) was reached. The probe was then kept at this position (in the X-Y dimension) at the given force offset. Then, negative pressure in the range between - 10 to -150 mbar was applied to aspirate cellular content. Before retracting the probe at the end of the aspiration process, the pressure was set back to 0 mbar. The force setpoint was adjusted by analyzing force distance curves from neighboring cells within the same experiment; the force value at which the curve takes a linear shape (in this case 80 nN) was estimated. This force value marks the point at which the probe makes contact with the glass bottom below the cell; consequently, this value was chosen as a setpoint for the extraction. Extraction of the ER fraction:

The experiments were performed using pyramidal cantilevers featuring an aperture area of 0.5 µm^2^ (see Fig. 1B) at -20 mbar using U2OS - or COS7 cells. Experiments for both cell lines were repeated twice on different days. Extraction of mitochondria and mitochondria pulling experiments were performed using pyramidal cantilevers featuring an aperture area of 1 µm^2^ or cantilevers with a cylindrical apex featuring an aperture area of 1.5 µm^2^ (see Fig. 1B) at -20 to -100 mbar using U2OS - or HeLa cells. Experiments including extraction of mitochondria were executed in 32 individual experimental setups on 32 individual days. Experiments showing recruitment of Drp1 were repeated four times on four different days with at least five cells per experiment in U2OS cells. Mitochondrial pulling experiments connected with calcium-imaging and thapsigargin-treatment (Fig. 5) were executed three times, each time on an individual day with at least seven cells per experiment.

For probe insertion with minimal Ca^2+^ influx as shown in Movie S7, newly coated (Sigmacote, see above) FluidFM cantilevers were used. To deliberately disturb the cell membrane (Movie S9), the probe was driven into the cell using the same setpoint, then the optical table (Newport) was gently flicked with the index finger to cause the probe to shift slightly, effectively disturbing the cell membrane. Cell viability was controlled using the LIVE/DEAD cell imaging kit (ThermoFisher).

### Mitochondrial transplantation experiments

FluidFM injection of bulk-purified mitochondria: Mitochondria were purified from approximately 2*10^6^ cultured HeLa cells continuously expressing the mitochondrial matrix marker su9-mCherry using the Qproteome Mitochondria Isolation Kit (Qiagen) following manufacturer’s instructions. After purification, the mitochondria were washed two times in injection buffer (see above), and finally resuspended in 40 µl injection buffer before being loaded into FluidFM-probes having a sharpened cylindrical apex.

Coupling mitochondrial extraction with transplantation from individual cell to cell: Mitochondria were aspired as described above, using FluidFM-probes having a sharpened cylindrical apex from HeLa-cells that were co-cultured on the same dish, previously seeded within another quadrant of the 4-well micro insets. Subsequently, the probe was moved to a region containing cells targeted for transplantation. For injection of mitochondria, the probe was positioned above a target cell as described above for the extraction of organelles. The cantilever containing the organelles was then inserted into the cell in contact mode (setpoint 100 nN). The injection process is controlled by observing the process in brightfield in real-time, at first, a positive pressure of 20 mbar was applied and the separation line between extract and perfluorooctane was monitored, the pressure was increased until the line was moving towards the aperture (maximal value of 200 mbar), effectively pushing the cell extract into the recipient cell. When the extract was totally or in large parts transferred to the recipient cell, the pressure was then set to 0 mbar and the probe was retracted. Z-stacks were taken to control for successfully transferred mitochondria. Cell viability was controlled using the LIVE/DEAD cell imaging kit (ThermoFisher) following manufacturers instruction. Both experimental approaches were performed in six individual experiments, each on individual days.

### Quantification of genetic incorporation of transplanted mitochondria

All experiments were conducted in biological triplicates. (A) FluidFM-injection of purified mitochondria: Single HeLa cells continuously expressing the mitochondrial matrix marker su9-BFP were seeded into a single quadrant of a 4-well micro-insert (ibidi) 72h before the experiment, at the day of the experiment, each seeded well contained 4-6 cells. All cells were injected as described above. (B) Coupling mitochondrial extraction with transplantation from individual cell to cell: One quadrant of a 4-well micro-insert (ibidi) was seeded with recipient HeLa (su9-BFP) cells as described above. A second quadrant was seeded with U2OS cells continuously expressing su9-mCherry as donors. The transplantation was executed as described above. Subsequently the U2OS cells were removed, again using a FluidFM probe filled with 1% sodium dodecyl sulfate (Merck). Before sequencing, the whole cell population was controlled for any signal mCherry positive-cells via fluorescence activated cell sorting, but no unwanted carryover of remaining U2OS cells (mCherry+) could be detected. Control for non-specific mitochondrial carryover: Approximately 100 000 Hela cells expressing the mitochondrial matrix marker su9-BFP were seeded onto a culture dish. They were mixed with purified mitochondria from approximately 2*10^6^ U2OS cells expressing su9-mCherry by pipetting the extracts gently on top of the cultured cells. Approaches A and B were grown for 8 days, transferred into a new culture dish and grown for another 6 days. Approach C was grown for 3 days, transferred into a new culture dish and grown for another 3 days. Subsequently, the total DNA of all samples was purified using the MasterPure™ Complete DNA and RNA Purification Kit (epicentre) following manufacturer’s instructions. Part of the D-loop region was amplified via PCR using primers 1 & 2 (table S2). To create a ‘null hypothesis’ for the following sequencing workflow, three test-samples were created consisting of 0.1% / 0.5% / 1% U2OS amplicons, mixed with HeLa amplicons. Subsequently all samples were sequenced using the a Pacbio Sequel SMRT cell, subsequently analyzed as described in Russo et al.(*68*) The reads were assigned to either U2OS mtDNA, or Hela mtDNA using four conserved mutations of U2OS cells within this region previously identified via Sanger-Sequencing (15959G>T, 16069C>T, 16108C>T, 16126T>C; see Table S1).

### Statistics and reproducibility

All representative experiments were repeated at least three times independently with similar results. Absolute counts of cells or mitochondrial particles are included in the main text or in the figure legends. Analysis for single nucleotide polymorphisms was performed as described in Russo et al. (68). Performed statistical tests and significance levels are indicated in the figures and respective figure legends.

### Analysis of mitochondrial quality control mechanisms with mitochondrial transplantation in primary HEKa cells

Cells were seeded into as described above, into low µ-dishes (ibidi) inside two-well culture inserts (ibidi). The special separation allows for drug-treatments of the donor cell population before the experiment, while the host cell population of HEKa cells remains under standard culture conditions. At the beginning of the experiment, the cell media was replaced with CO_2_-independent growth medium containing 10% FBS (ThermoFisher) and 1% penicillin-streptomycin (ThermoFisher). 4 µM Oligomycin was added when indicated. Treatment conditions: 10 µM CCCP for 3 h, 8 µM Doxycycline for 48 h, 750 µM H_2_O_2_ for 3 h. Mitochondria were extracted from Hela cells and transplanted into HEKa cells. A Z-stack image series of the transplant and the host cell was taken directly after the transplantation process with a 63× oil objective with 1.4 numerical aperture and a 2× lens switcher in 500 nm steps. Further Z-stacks were taken at other time-points depending on the individual experiment. For the endpoint at 18 - 22 h post transplantation, the mitochondrial network was visualized using MitoTracker® Green FM (ThermoFisher) following manufacturers instructions. The overlap between the transplant and the host mitochondrial network was controlled. For the quantitative analysis, data was analyzed with self-written Matlab (R2018a) scripts.

### Split-kinesin experiments

pKIF5C-HA-FRB was a kind gift from Prof. Sean Munro. Expresses fusion of Rat kinesin minus tail to FRB; pFKBP-mCh-Fis1TM was subcloned from pFKBP-GFP-myc-GRIP (Sean Munro) by swapping the GFP-myc-GRIP fragment with mCherry fused to the TM domain of yeast Fis1 for outer-mitochondrial membrane targeting. U2OS cells stably expressing mtBFP and a shRNA against DRP1 (53) were reverse transfected with the two constructs above. 2 days later cells were imaged using a spinning disk microscope. Rapamycin was diluted in growth medium to a final concentration of 1 nM. A microfluidic imaging device was used to allow (rapamycin-containing) medium replacement while imaging. Experiments were performed twice on different days.

## Acknowledgements

We thank Maximilian Mittelviefhaus (ETH Zurich), Patrick Frederix (Nanosurf AG, Switzerland), Pablo Dörig, and Dario Ossola (Cytosurge AG, Switzerland) for their support, as well as Stephen Wheeler (ETH Zurich) and the personnel of ETH ScopeM facility and the Functional Genomics Center Zurich for technical assistance. We thank Sean Munro for plasmids. This work was supported by a grant from the Volkswagen foundation (Initiative “Life” to J.A.V.), a European Research Council Advanced Grant (no. 883077 to J.A.V.), a European Research Commission Starting Grant (no. 337906 to B.K.), and the EUROSTARS project grant E!11644 “SOUL” (to E.S. and T.Z.).

## Author contributions

C.G.G., B.K. and J.A.V designed the study. C.G.G. and J.A.V. wrote the manuscript with input from all authors. C.G.G. and Q.F. designed, executed and analyzed the experiments. C.G.G., O.G.G., E.S. and T.Z. designed and manufactured FluidFM-probes.

## Supplemental Material

Supplementary text ‘Force-induced mitochondrial fission’, Supplementary Fig. 1-13; Supplementary Movies 1-12, Supplementary Tables 1 & 2

## Data availability

Data to understand and assess the conclusion of this research are available in the main text, raw data sets will be made available upon request.

## Code availability

Matlab code used for image analysis and mitochondrial particle distribution will be made available upon request.

## References

1. C. Wang, R. J. Youle, The role of mitochondria in apoptosis. Annu. Rev. Genet. 43, 95–118 (2009).

2. A. P. West, W. Khoury-Hanold, M. Staron, M. C. Tal, C. M. Pineda, S. M. Lang, M. Bestwick, B. A. Duguay, N. Raimundo, D. A. MacDuff, S. M. Kaech, J. R. Smiley, R. E. Means, A. Iwasaki, G. S. Shadel, Mitochondrial DNA stress primes the antiviral innate immune response. Nature. 520, 553–557 (2015).

3. Z. Wu, S. Oeck, A. P. West, K. C. Mangalhara, A. G. Sainz, L. E. Newman, X. O. Zhang, L. Wu, Q. Yan, M. Bosenberg, Y. Liu, P. L. Sulkowski, V. Tripple, S. M. Kaech, P. M. Glazer, G. S. Shadel, Mitochondrial DNA stress signalling protects the nuclear genome. Nat. Metab. 1, 1209–1218 (2019).

4. R. P. Chakrabarty, N. S. Chandel, Mitochondria as Signaling Organelles Control Mammalian Stem Cell Fate. Cell Stem Cell. 28, 394–408 (2021).

5. N. Sun, R. J. Youle, T. Finkel, The Mitochondrial Basis of Aging. Mol. Cell. 61, 654–666 (2016).

6. T. E. S. Kauppila, J. H. K. Kauppila, N. G. Larsson, Mammalian Mitochondria and Aging: An Update. Cell Metab. 25, 57–71 (2017).

7. S. L. Archer, Mitochondrial dynamics - Mitochondrial fission and fusion in human diseases. N. Engl. J. Med. 369, 2236–2251 (2013).

8. P. Mishra, D. C. Chan, Mitochondrial dynamics and inheritance during cell division, development and disease. Nat. Rev. Mol. Cell Biol. 15, 634–646 (2014).

9. J. D. Vevea, T. C. Swayne, I. R. Boldogh, L. A. Pon, Inheritance of the fittest mitochondria in yeast. Trends Cell Biol. 24, 53–60 (2014).

10. M. A. Clark, J. W. Shay, Mitochondrial transformation of mammalian cells. Nature. 295, 605–607 (1982).

11. D. Liu, Y. Gao, J. Liu, Y. Huang, J. Yin, Y. Feng, L. Shi, B. P. Meloni, C. Zhang, M. Zheng, J. Gao, Intercellular mitochondrial transfer as a means of tissue revitalization. Signal Transduct. Target. Ther. 6 (2021), doi:10.1038/s41392-020-00440-z.

12. T. Santos-Ferreira, S. Llonch, O. Borsch, K. Postel, J. Haas, M. Ader, Retinal transplantation of photoreceptors results in donor-host cytoplasmic exchange. Nat. Commun. 7, 1–7 (2016).

13. A. Rustom, R. Saffrich, I. Markovic, P. Walther, H. H. Gerdes, Nanotubular Highways for Intercellular Organelle Transport. Science 303, 1007–1010 (2004).

14. A. Caicedo, P. M. Aponte, F. Cabrera, C. Hidalgo, M. Khoury, Artificial mitochondria transfer: current challenges, Advances, and Future Applications. Stem Cells Int. 2017, 7610414 (2017).

15. B. V. Chernyak, Mitochondrial Transplantation: A Critical Analysis. Biochem. 85, 636–641 (2020).

16. P. Actis, Sampling from Single Cells. small. 1700300, 1–11 (2018).

17. B. P. Nadappuram, P. Cadinu, A. Barik, A. J. Ainscough, M. J. Devine, M. Kang, J. Gonzalez-Garcia, J. T. Kittler, K. R. Willison, R. Vilar, P. Actis, B. Wojciak-Stothard, S. H. Oh, A. P. Ivanov, J. B. Edel, Nanoscale tweezers for single-cell biopsies. Nat. Nanotechnol. 14, 80–88 (2019).

18. E. N. Tóth, A. Lohith, M. Mondal, J. Guo, A. Fukamizu, N. Pourmand, Single-cell nanobiopsy reveals compartmentalization of mRNAs within neuronal cells. J. Biol. Chem. 293, 4940–4951 (2018).

19. O. Guillaume-Gentil, T. Rey, P. Kiefer, A. J. Ibáñez, R. Steinhoff, R. Brönnimann, L. Dorwling-Carter, T. Zambelli, R. Zenobi, J. A. Vorholt, Single-Cell Mass Spectrometry of Metabolites Extracted from Live Cells by Fluidic Force Microscopy. Anal. Chem. 89, 5017–5023 (2017).

20. H. Zhu, Q. Li, T. Liao, X. Yin, Q. Chen, Z. Wang, M. Dai, L. Yi, S. Ge, C. Miao, W. Zeng, L. Qu, Z. Ju, G. Huang, C. Cang, W. Xiong, Metabolomic profiling of single enlarged lysosomes. Nat. Methods. 18, 788–798 (2021).

21. T. Wu, T. Teslaa, S. Kalim, C. T. French, S. Moghadam, R. Wall, F. Miller, X. O. N. Witte, M. A. Teitell, P. Chiou, Photothermal Nanoblade for Large Cargo Delivery into Mammalian. Anal. Chem. 83, 1321–1327 (2011).

22. T. Wu, E. Sagullo, D. Case, T. G. Graeber, P. Chiou, M. A. Teitell, T. Wu, E. Sagullo, D. Case, X. Zheng, Y. Li, J. S. Hong, T. Teslaa, Mitochondrial Transfer by Photothermal Nanoblade Restores Metabolite Profile in Mammalian Cells. Cell Metab. 23, 921–929 (2016).

23. A. Meister, M. Gabi, P. Behr, P. Studer, J. Vörös, P. Niedermann, J. Bitterli, J. Polesel-Maris, M. Liley, H. Heinzelmann, T. Zambelli, FluidFM: Combining atomic force microscopy and nanofluidics in a universal liquid delivery system for single cell applications and beyond. Nano Lett. 9, 2501–2507 (2009).

24. O. Guillaume-Gentil, E. Potthoff, D. Ossola, C. M. Franz, T. Zambelli, J. A. Vorholt, Force-controlled manipulation of single cells: From AFM to FluidFM. Trends Biotechnol. 32, 381–388 (2014).

25. O. Guillaume-Gentil, E. Potthoff, D. Ossola, P. Dörig, T. Zambelli, J. A. Vorholt, Force-controlled fluidic injection into single cell nuclei. Small. 9, 1904–1907 (2013).

26. O. Guillaume-Gentil, R. V. V. Grindberg, R. Kooger, L. Dorwling-Carter, V. Martinez, D. Ossola, M. Pilhofer, T. Zambelli, J. A. A. Vorholt, Tunable Single-Cell Extraction for Molecular Analyses. Cell. 166, 506–517 (2016).

27. R. Milo, P. Jorgensen, U. Moran, G. Weber, M. Springer, BioNumbers The database of key numbers in molecular and cell biology. Nucleic Acids Res. 38, 750–753 (2009).

28. J. Wu, L. Liu, T. Matsuda, Y. Zhao, A. Rebane, M. Drobizhev, Y. F. Chang, S. Araki, Y. Arai, K. March, T. E. Hughes, K. Sagou, T. Miyata, T. Nagai, W. H. Li, R. E. Campbell, Improved orange and red Ca2+ indicators and photophysical considerations for optogenetic applications. ACS Chem. Neurosci. 4, 963–972 (2013).

29. J. B. Barbara Stolz, Sequestration of iontophoretically injected calcium by living endothelial cells. Cell Calcium. 8, 103–121 (1987).

30. D. Gonzalez-Rodriguez, S. Sart, A. Babataheri, D. Tareste, A. I. Barakat, C. Clanet, J. Husson, Elastocapillary Instability in Mitochondrial Fission. Phys. Rev. Lett. 115, 1–5 (2015).

31. E. Smirnova, D. L. Shurland, S. N. Ryazantsev, A. M. Van Der Bliek, A human dynamin-related protein controls the distribution of mitochondria. J. Cell Biol. 143, 351–358 (1998).

32. D. A. Rendon, Important methodological aspects that should be taken into account during the research of isolated mitochondria. Anal. Biochem. 589, 113492 (2020).

33. S. R. Denison, F. Wang, N. A. Becker, B. Schüle, N. Kock, L. A. Phillips, C. Klein, D. I. Smith, Alterations in the common fragile site gene Parkin in ovarian and other cancers. Oncogene. 22, 8370–8378 (2003).

34. J. Gao, A. Qin, D. Liu, R. Ruan, Q. Wang, J. Yuan, T. S. Cheng, A. Filipovska, J. M. Papadimitriou, K. Dai, Q. Jiang, X. Gao, J. Q. Feng, H. Takayanagi, C. Zhang, M. H. Zheng, Endoplasmic reticulum mediates mitochondrial transfer within the osteocyte dendritic network. Sci. Adv. 5, 1–13 (2019).

35. J. R. Brestoff, C. B. Wilen, J. R. Moley, Y. Li, W. Zou, N. P. Malvin, M. N. Rowen, B. T. Saunders, H. Ma, M. R. Mack, B. L. Hykes, D. R. Balce, A. Orvedahl, J. W. Williams, N. Rohatgi, X. Wang, M. R. McAllaster, S. A. Handley, B. S. Kim, J. G. Doench, B. H. Zinselmeyer, M. S. Diamond, H. W. Virgin, A. E. Gelman, S. L. Teitelbaum, Intercellular Mitochondria Transfer to Macrophages Regulates White Adipose Tissue Homeostasis and Is Impaired in Obesity. Cell Metab. 33, 270-282.e8 (2021).

36. K. Hayakawa, E. Esposito, X. Wang, Y. Terasaki, Y. Liu, C. Xing, X. Ji, E. H. Lo, Transfer of mitochondria from astrocytes to neurons after stroke. Nature. 535, 551–555 (2016).

37. M. Yaar, B. A. Gilchrest, Ageing and photoageing of keratinocytes and melanocytes. Clin. Exp. Dermatol. 26, 583–591 (2001).

38. A. M. van der Bliek, Q. Shen, S. Kawajiri, Mechanisms of mitochondrial fission and fusion. Cold Spring Harb. Perspect. Biol. 5, a011072 (2013).

39. M. Zachari, N. T. Ktistakis, Mammalian Mitophagosome Formation: A Focus on the Early Signals and Steps. Front. Cell Dev. Biol. 8, 1–11 (2020).

40. D. Narendra, A. Tanaka, D. F. Suen, R. J. Youle, Parkin is recruited selectively to impaired mitochondria and promotes their autophagy. J. Cell Biol. 183, 795–803 (2008).

41. N. Matsuda, S. Sato, K. Shiba, K. Okatsu, K. Saisho, C. A. Gautier, Y. S. Sou, S. Saiki, S. Kawajiri, F. Sato, M. Kimura, M. Komatsu, N. Hattori, K. Tanaka, PINK1 stabilized by mitochondrial depolarization recruits Parkin to damaged mitochondria and activates latent Parkin for mitophagy. J. Cell Biol. 189, 211–221 (2010).

42. N. Moullan, L. Mouchiroud, X. Wang, D. Ryu, E. G. Williams, A. Mottis, V. Jovaisaite, M. V. Frochaux, P. M. Quiros, B. Deplancke, R. H. Houtkooper, J. Auwerx, Tetracyclines disturb mitochondrial function across eukaryotic models: A call for caution in biomedical research. Cell Rep. 10, 1681–1691 (2015).

43. M. Yang, L. Luna, J. G. Sørbø, I. Alseth, R. F. Johansen, P. H. Backe, N. C. Danbolt, L. Eide, M. Bjørås, Human OXR1 maintains mitochondrial DNA integrity and counteracts hydrogen peroxide-induced oxidative stress by regulating antioxidant pathways involving p21. Free Radic. Biol. Med. 77, 41–48 (2014).

44. A. J. Sercel, A. N. Patananan, T. Man, T. H. Wu, A. K. Yu, G. W. Guyot, S. Rabizadeh, K. R. Niazi, P. Y. Chiou, M. A. Teitell, Stable transplantation of human mitochondrial DNA by high-throughput, pressurized isolated mitochondrial delivery. Elife. 10, 1–45 (2021).

45. Y.-W. Yang, M. D. Koob, Transferring isolated mitochondria into tissue culture cells. Nucleic Acids Res. 40, e148 (2012).

46. Y. G. Yoon, M. D. Koob, Intramitochondrial transfer and engineering of mammalian mitochondrial genomes in yeast. Mitochondrion. 46, 15–21 (2019).

47. L. S. Kaguni, DNA polymerase γ, the mitochondrial replicase. Annu. Rev. Biochem. 73, 293–320 (2004).

48. M. T. Lott, J. N. Leipzig, O. Derbeneva, H. Michael Xie, D. Chalkia, M. Sarmady, V. Procaccio, D. C. Wallace, MtDNA variation and analysis using Mitomap and Mitomaster. Curr. Protoc. Bioinforma., 1–26 (2013).

49. D. Bogenhagen, D. A. Clayton, The number of mitochondrial deoxyribonucleic acid genomes in mouse L and human HeLa cells. Quantitative isolation of mitochondrial deoxyribonucleic acid. J. Biol. Chem. 249, 7991–7995 (1974).

50. M. Satoh, T. Kuroiwa, Organization of multiple nucleoids and DNA molecules in mitochondria of a human cell. Exp. Cell Res. 196, 137–140 (1991).

51. C. Kukat, C. A. Wurm, H. Spåhr, M. Falkenberg, N. G. Larsson, S. Jakobs, Super-resolution microscopy reveals that mammalian mitochondrial nucleoids have a uniform size and frequently contain a single copy of mtDNA. Proc. Natl. Acad. Sci. U. S. A. 108, 13534–13539 (2011).

52. W. Chen, S. Li, A. S. Kulkarni, L. Huang, J. Cao, K. Qian, J. Wan, Single Cell Omics: From Assay Design to Biomedical Application. Biotechnol. J. 15, 1–10 (2020).

53. S. C. J. Helle, Q. Feng, M. J. Aebersold, L. Hirt, R. R. Grüter, A. Vahid, A. Sirianni, S. Mostowy, J. G. Snedeker, A. Šarić, T. Idema, T. Zambelli, B. Kornmann, Mechanical force induces mitochondrial fission. Elife. 6, 1–26 (2017).

54. J. R. Friedman, L. L. Lackner, M. West, J. R. DiBenedetto, J. Nunnari, G. K. Voeltz, ER tubules mark sites of mitochondrial division. Science 334, 358–362 (2011).

55. S. Jung, C. Clanet, J. W. M. Bush, Capillary instability on an elastic helix. Soft Matter. 10, 3225–3228 (2014).

56. Q. S. Li, G. Y. H. Lee, C. N. Ong, C. T. Lim, AFM indentation study of breast cancer cells. Biochem. Biophys. Res. Commun. 374, 609–613 (2008).

57. V. J. J. Cadete, S. Deschênes, A. Cuillerier, F. Brisebois, A. Sugiura, A. Vincent, D. Turnbull, M. Picard, H. M. McBride, Y. Burelle, Formation of mitochondrial-derived vesicles is an active and physiologically relevant mitochondrial quality control process in the cardiac system. J. Physiol. 594, 5343–5362 (2016).

58. A. Sugiura, G. McLelland, E. A. Fon, H. M. McBride, A new pathway for mitochondrial quality control: mitochondrial-derived vesicles. EMBO J. 33, 2142–2156 (2014).

59. P. Katajisto, J. Döhla, C. L. Chaffer, N. Pentinmikko, N. Marjanovic, S. Iqbal, R. Zoncu, W. Chen, R. Weinberg, D. M. Sabatini, Asymmetric apportioning of aged mitochondria between daughter cells is required for stemness. Science 348, 340–343 (2015).

60. A. S. Moore, S. M. Coscia, C. L. Simpson, F. E. Ortega, E. C. Wait, J. M. Heddleston, J. J. Nirschl, C. J. Obara, P. Guedes-Dias, C. A. Boecker, T.-L. Chew, J. A. Theriot, J. Lippincott-Schwartz, E. L. F. Holzbaur, Actin cables and comet tails organize mitochondrial networks in mitosis. Nature 591. 659-664 (2021).

61. H. Mobarak, M. Heidarpour, P. S. J. Tsai, A. Rezabakhsh, R. Rahbarghazi, M. Nouri, M. Mahdipour, Autologous mitochondrial microinjection; A strategy to improve the oocyte quality and subsequent reproductive outcome during aging. Cell Biosci. 9, 1–15 (2019).

62. A. S. Reznichenko, C. Huyser, M. S. Pepper, Mitochondrial transfer: Implications for assisted reproductive technologies. Appl. Transl. Genomics. 11, 40–47 (2016).

63. V. I. Floros, A. Pyle, S. Dietmann, W. Wei, W. W. C. Tang, N. Irie, B. Payne, A. Capalbo, L. Noli, J. Coxhead, G. Hudson, M. Crosier, H. Strahl, Y. Khalaf, M. Saitou, D. Ilic, M. A. Surani, P. F. Chinnery, Segregation of mitochondrial DNA heteroplasmy through a developmental genetic bottleneck in human embryos. Nat. Cell Biol. 20, 144–151 (2018).

64. T. Lieber, S. P. Jeedigunta, J. M. Palozzi, R. Lehmann, T. R. Hurd, Mitochondrial fragmentation drives selective removal of deleterious mtDNA in the germline. Nature. 570, 380–384 (2019).

65. L. S. Ludwig, C. A. Lareau, J. C. Ulirsch, E. Christian, C. Muus, L. H. Li, K. Pelka, W. Ge, Y. Oren Brack, T. Law, C. Rodman, J. H. Chen, G. M. Boland, N. Hacohen, O. Rozenblatt-Rosen, M. J. Aryee, J. D. Buenrostro, A. Regev, V. G. Sankaran, Lineage Tracing in Humans Enabled by Mitochondrial Mutations and Single-Cell Genomics. Cell. 176, 1325-1339.e22 (2019).

66. J. Schindelin, I. Arganda-Carreras, E. Frise, V. Kaynig, M. Longair, T. Pietzsch, S. Preibisch, C. Rueden, S. Saalfeld, B. Schmid, J.-Y. Tinevez, D. J. White, V. Hartenstein, K. Eliceiri, P. Tomancak, Cardona, Fiji: an open-source platform for biological-image analysis. Nat. Methods. 9, 676–82 (2012).

67. K. M. Thyng, C. A. Greene, R. D. Hetland, H. M. Zimmerle, S. F. DiMarco, True colors of oceanography. Oceanography. 29, 9–13 (2016).

68. G. Russo, A. Patrignani, L. Poveda, F. Hoehn, B. Scholtka, R. Schlapbach, A. M. Garvin, Highly sensitive, non-invasive detection of colorectal cancer mutations using single molecule, third generation sequencing. Appl. Transl. Genomics. 7, 32–39 (2015).

