## Supplementary Materials for "Mitochondria transplantation between living cells"

**This PDF file includes:**

Supplementary Text on mitochondrial fission
Fig. S1 to S13
Captions for Movies 1 to 12
Tables 1 & 2

**Other Supplementary Materials for this manuscript include the following:**
Captions for Supplementary Movies 1 to 12

**Supplementary Text**

Force-induced mitochondrial fission

It has previously been suggested that mitochondrial membrane constriction is a prerequisite for mitochondria fission (1, 2); however it was impossible to exert highly localized hydrodynamic pulling forces intracellularly with sub micrometer resolution. FluidFM has the advantage of allowing to distinguish between mechanical force exertion and other cellular processes possibly involved such as calcium signaling. When extracting mitochondria, we observed induction of the pearls-on-a-string phenotype on mitochondria, followed by division of the inner-and outer mitochondrial membrane (Supplementary Fig. 5A,B, Supplementary Movies 5 and 6). We wondered whether the scission process was due to mechanical forces exerted by FluidFM or by recruitment of the native mitochondrial fission machinery to these sites. A main component of this machinery is Drp1, a mechanoenzyme that assembles circularily around mitochondria and uses the energy from GTP hydrolysis to mediate membrane scission (3). To assess the recruitment of the fission machinery to constricted sites, we expressed a fluorescently labelled version of Drp1 (Drp1-mCherry) in U2OS cells and performed mitochondrial pulling experiments to follow the fluorescent signals of both the mitochondrial matrix and Drp1. We observed that the FluidFM-induced pearls-on-a-string phenotype led to the recruitment of fluorescently labelled Drp1 (Drp1-mCherry) at the induced constriction sites of targeted mitochondrial tubes (Supplementary Fig. 5D, n = 18), thus providing a direct link between induced constriction and recruitment of Drp1.

The observed force-induced shape transition leads to the question of its relevance *in vivo*. To investigate this question further we examined kinesin as an endogenous motor protein (4) as a potential trigger. We employed the split-kinesin strategy using rapamycin-inducible protein interactions (5). Briefly, FK506 binding protein (FKBP) was fused to mCherry and to the transmembrane domain of Fis1 for OMM targeting. Its partner, FKBP–rapamycin binding (FRB), was fused 20 to the motor domain of kinesin. Upon rapamycin exposure, FKBP and FRB form a stable complex, thereby coupling mitochondria to the kinesin motor domain and directs their transport on microtubules. Addition of rapamycin to cells expressing these constructs induced a global shape transition of mitochondria similar to that observed upon FluidFM aspiration (Supplementary Fig. 5E), suggesting that hydrodynamic pulling forces created by FluidFM aspiration are in a similar range as forces created by kinesin motor proteins.

These results are congruent with previous studies suggesting that mitochondrial membrane constriction is a prerequisite for mitochondria fission (1, 2, 6). An indicator of plasma membrane damage and an inducer of mitochondrial pearling is leakage of calcium ions into the cytoplasm and mitochondria (7). Mammalian cells tightly control calcium concentrations whereby the ER acts as $Ca^{2+}$ storage compartment and mitochondrial pearling and subsequent fission has previously been associated with $Ca^{2+}$ signaling (8, 9). To investigate whether $Ca^{2+}$ flux is associated with shape transition of mitochondria in our approach, we used the $Ca^{2+}$ sensitive fluorophore mito-R-GECO1 (10) targeted to the mitochondrial matrix and followed mitochondrial $Ca^{2+}$ dynamics in time-lapse microscopy experiments. We observed no change in signal intensity of R-GECO1, neither after probe insertion, nor during the extraction process (Supplementary Movie 7). To control for functionality of the sensor system, we probed the initially extracted cells twice (n =17). In the second approach, we manually displaced the probe while it was inside the cell, inducing rupture of the cytoplasmic membrane. Because the cell culture medium contains roughly a 6000-fold excess of calcium compared to mitochondria (7), we expected an influx of $Ca^{2+}$. Indeed, a systemic $Ca^{2+}$ influx signal of mito-R-GECO occurred, propagating radially from the probe insertion site, followed by rounding of mitochondria and cell death (Supplementary Movie 8), in line with $Ca^{2+}$ inducing apoptosis via cytochrome *c* release from mitochondria (7). A similar mitochondrial calcium influx was observed when using imperfectly coated probes that we expected to result in localized $Ca^{2+}$ influx upon membrane puncture. Indeed, we observed a rapid and transient increase of R-GECO1 fluorescence intensity in mitochondria upon probe insertion. However, there was no immediate influence on the mitochondrial morphology. Only when negative pressure was applied, 'pearling' of mitochondrial tubes was observed exclusively in direct proximity of the aperture. The morphology of mitochondria situated further away did not change, despite being equally affected by the $Ca^{2+}$ influx (n=14) (Supplementary Fig. 6A and Supplementary Movie 9). Finally, we depleted $Ca^{2+}$ in the medium by adding the chelating agent EGTA. Under this condition, the signal intensity of R-GECO1 did not rise, neither upon probe entry nor during the mechanically induced fission process (Supplementary Fig. 6B and Supplementary Movie 10). To rule out an involvement of calcium stored within the ER, we treated U2OS-cells with thapsigargin, which depletes the ER calcium reservoir. We did not detect any impact of thapsigargin on force induced pearling of mitochondria (Supplementary Fig. 6C). Based on these results, we conclude that calcium influx is not linked to the observed mitochondrial shape transition. Due to the directional propagation along mitochondrial tubes, rather than a radial propagation from the site of probe insertion, as well as lacking evidence of an influence of calcium on the observed process, we conclude that hydrodynamic pulling forces rather than a biochemical signal is responsible for mitochondrial pearling and fission.

1. L. Carlini, D. Mahecic, T. Kleele, A. Roux, S. Manley, Membrane bending energy and tension govern mitochondrial division. *BioRxiv*, 1–34 (2018).
2. S. C. J. Helle *et al.*, Mechanical force induces mitochondrial fission. *Elife*. **6**, 1–26 (2017).
3. E. Smirnova, D. L. Shurland, S. N. Ryazantsev, A. M. Van Der Bliek, A human dynamin-related protein controls the distribution of mitochondria. *J. Cell Biol.* **143**, 351–358 (1998).
4. E. Meyhofer, J. Howard, The force generated by a single kinesin molecule against an elastic load. *Proc. Natl. Acad. Sci.* **92**, 574–578 (1995).
5. P. van Bergeijk, C. C. Hoogenraad, L. C. Kapitein, Right Time, Right Place: Probing the Functions of Organelle Positioning. *Trends Cell Biol.* **26**, 121–134 (2016).
6. B. Cho *et al.*, Constriction of the mitochondrial inner compartment is a priming event for mitochondrial division. *Nat. Commun.* **8**, 15754 (2017).
7. L. Ghibelli, C. Cerella, M. Diederich, The dual role of calcium as messenger and stressor in cell damage, death, and survival. *Int. J. Cell Biol.* **2010** (2010), doi:10.1155/2010/546163.
8. A. M. van der Bliek, Q. Shen, S. Kawajiri, Mechanisms of mitochondrial fission and fusion. *Cold Spring Harb. Perspect. Biol.* **5** (2013), doi:10.1101/cshperspect.a011072.
9. N. Nemani *et al.*, MIRO-1 Determines Mitochondrial Shape Transition upon GPCR Activation and Ca2+ Stress. *Cell Rep.* **23**, 1005–1019 (2018).
10. J. Wu *et al.*, Improved orange and red Ca2+ indicators and photophysical considerations for optogenetic applications. *ACS Chem. Neurosci.* **4**, 963–972 (2013).

### Supplementary Figures

**A**

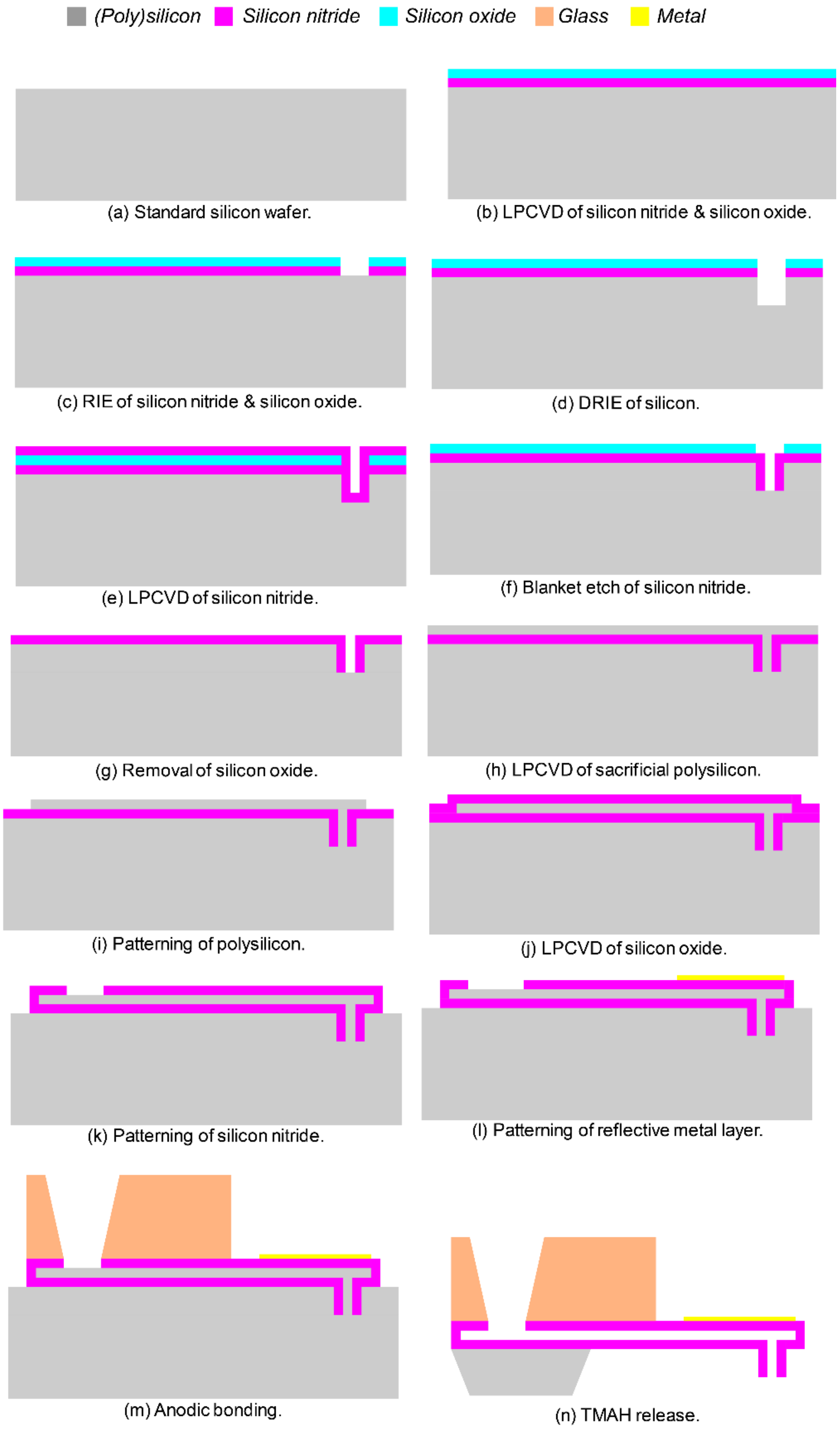

B

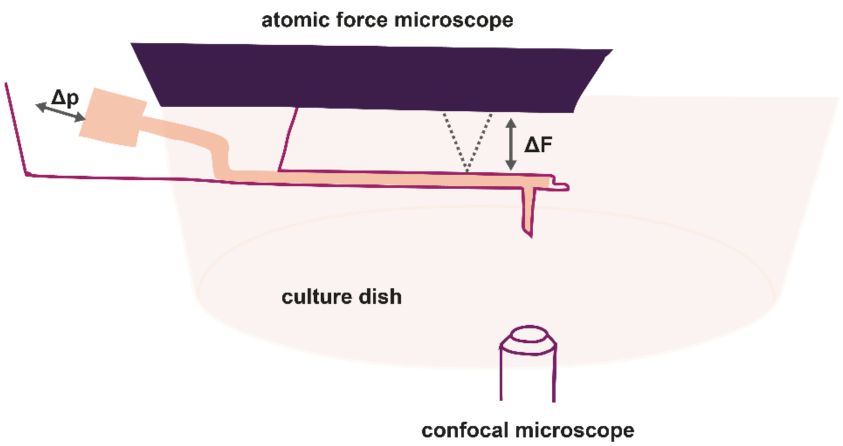

**Supplementary Fig. 1.**
**(A)** FluidFM fabrication process. (**a** - **n**) Steps for the fabrication of hollow FluidFM cantilevers comprising a cylindrical apex. (**B**) Schematic overview of the FluidFM setup.

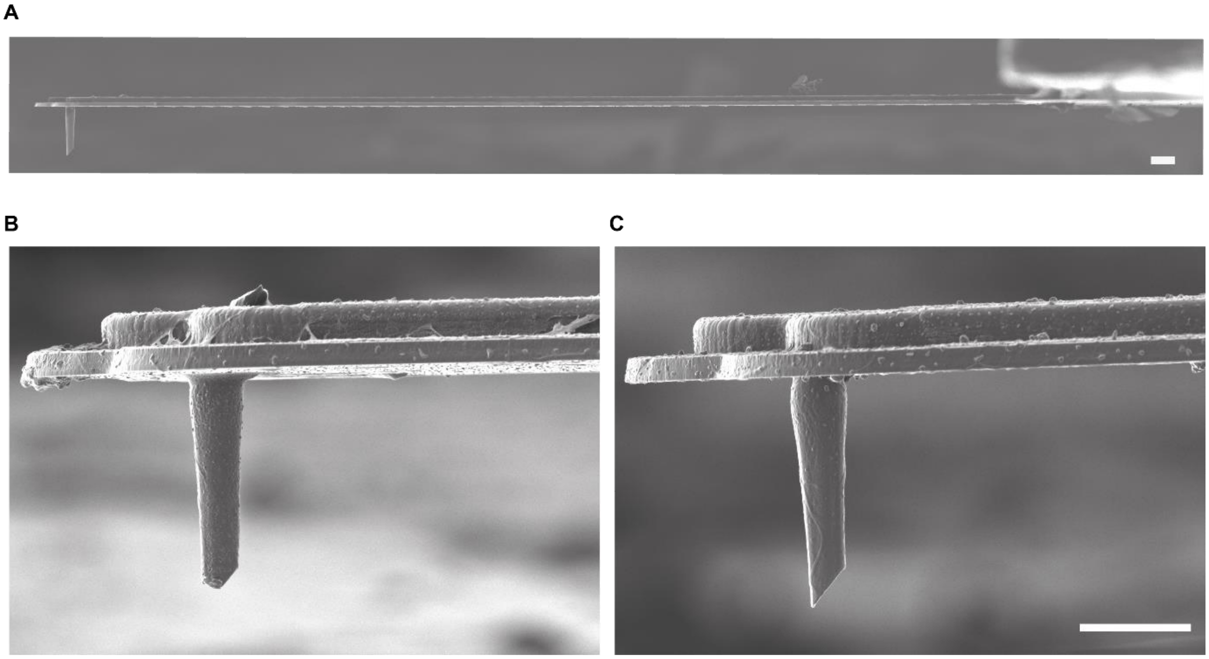

**Supplementary Fig. 2.**
Mechanical robustness of FluidFM cantilevers comprising a sharpened cylindrical apex. Focused Ion Beam images of FluidFM cantilevers: (**A**) Side view of a slanted cylindrical FluidFM cantilever with a 1.2 µm diameter of the cylinder. (**B**) Cantilever that was used for mitochondrial transplantation with a setpoint of 1000 nN, the apex is broken impairing insertion of the probe into cells (**C**) Cantilever that was used for mitochondrial transplantation with a setpoint of 400 nN, the apex remains intact. Scale bar: 10 µm

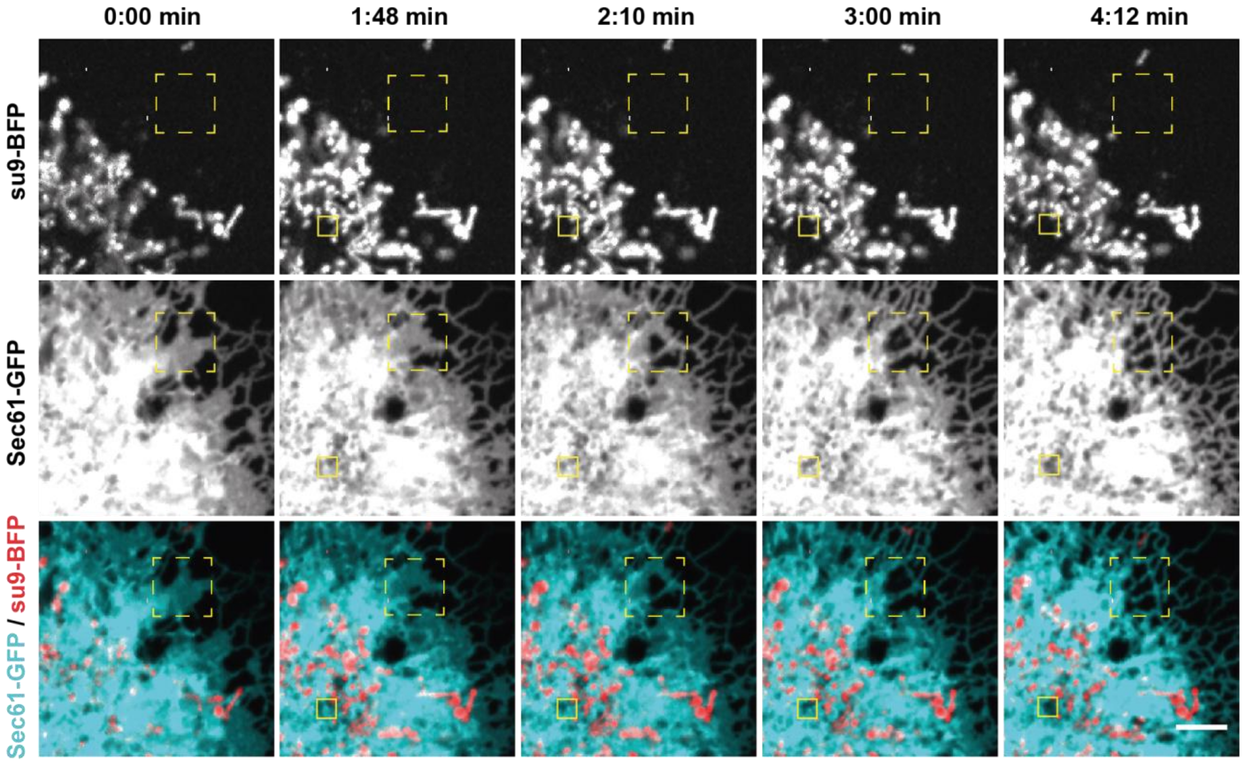

**Supplementary Fig. 3.**
ER extraction of COS7-cells. COS7-cells stably expressing the ER membrane marker Sec61-GFP and mitochondrial matrix marker su9-BFP. Small square indicates cantilever position, big dashed line indicates a zone of ER re-arrangement. Scale bar: 5 µm.

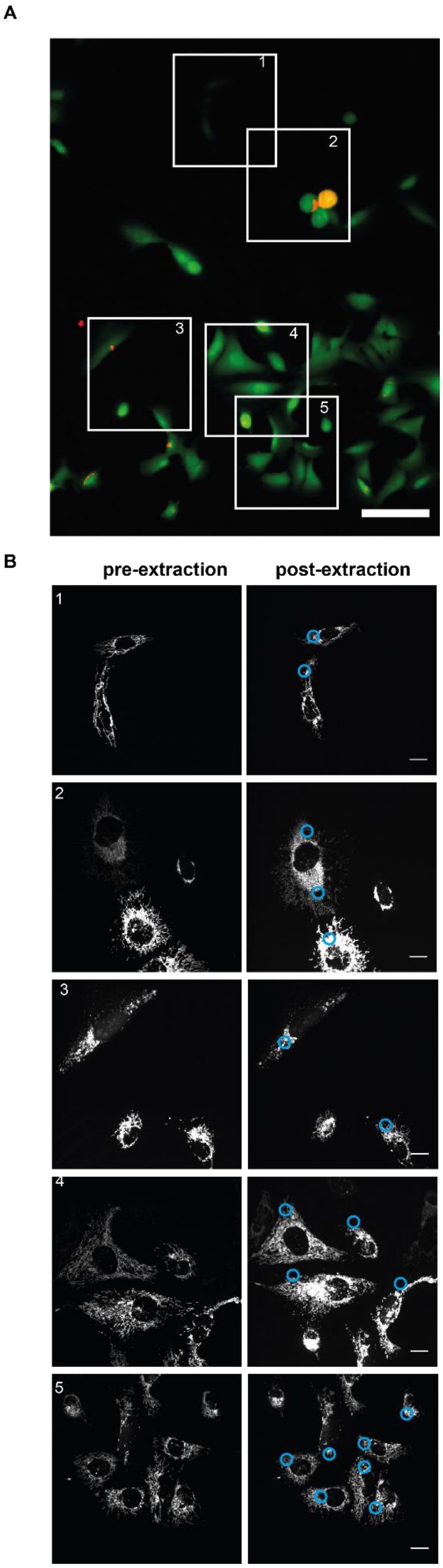

**Supplementary Fig. 4.**
Viability of Hela cells post extraction of mitochondria. (**A**) Overview of HeLa cells 2 h post mitochondrial extraction, stained with the LIVE-DEAD Cell imaging kit. Viable cells show green fluorescence signal, dead cells show red fluorescence. Areas 1 - 5 show regions with extracted cells. Scale bar: 50 µm. (**B**) Mitochondrial networks (su9-BFP) from the regions shown in a, before and 2 h post extraction. The insertion sites of the cantilevers for extraction are highlighted by blue circles. The experiment was conducted twice, 36 out of 37 sampled cells remained viable. Scale bar: 10 µm.

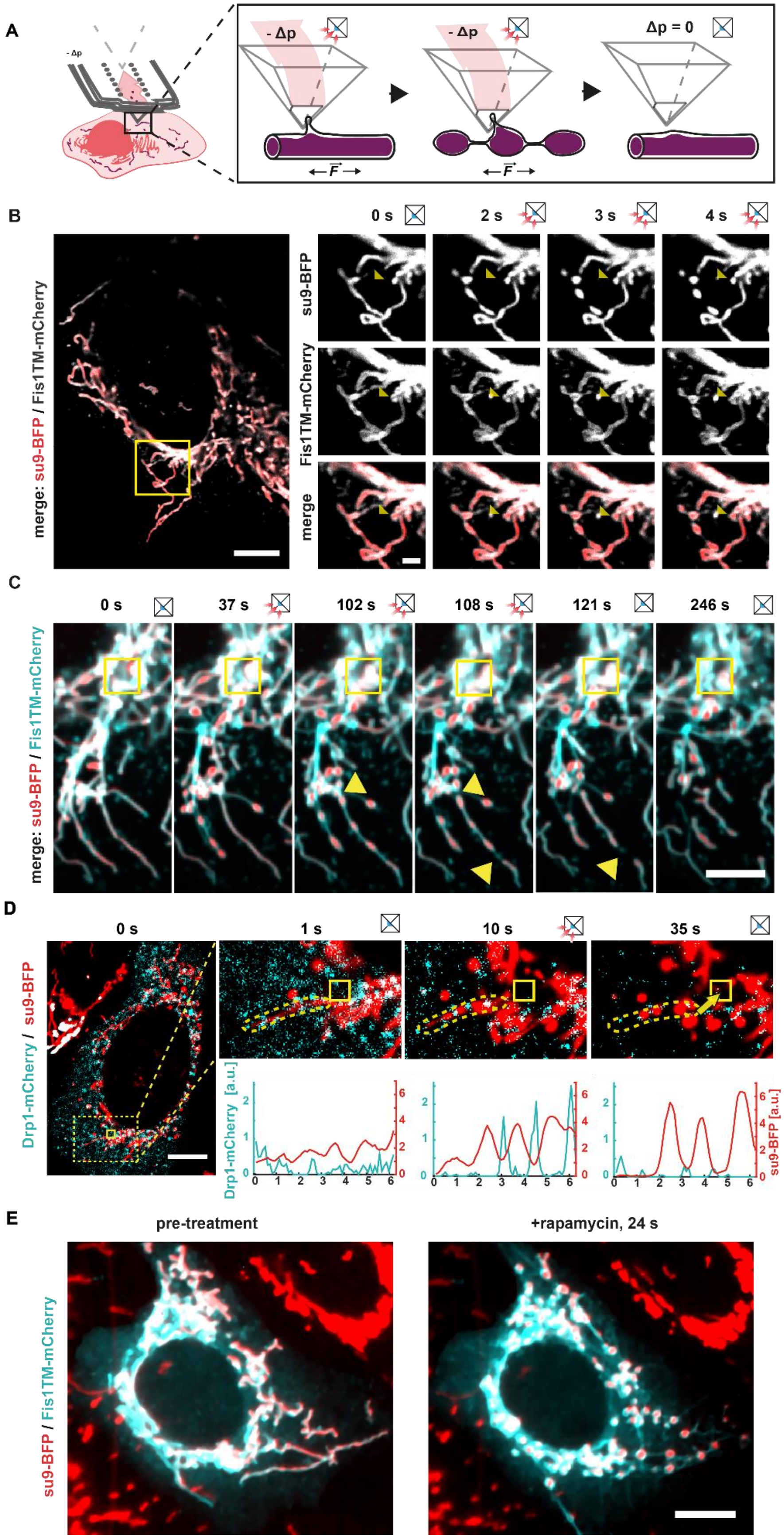

A
- Δp
- Δp
Δp = 0
B
0 s
2 s
3 s
4 s
merge: su9-BFP / Fis1TM-mCherry
su9-BFP
Fis1TM-mCherry
merge
C
0 s
37 s
102 s
108 s
121 s
246 s
merge: su9-BFP / Fis1TM-mCherry
D
0 s
1 s
10 s
35 s
Drp1-mCherry / su9-BFP
Drp1-mCherry [a.u.]
su9-BFP [a.u.]
E
pre-treatment
+rapamycin, 24 s
su9-BFP / Fis1TM-mCherry

**Supplementary Fig. 5.**

Mitochondrial shape transition and fission. **(A),** Schematic representation of the working model of the shape transition into the pearls-on-a-string phenotype upon exertion of pulling forces. Small box icons represent cantilever apexes applying negative hydrodynamic forces (-Δp), or without pressure differences (Δp = 0). (**B**) Images of U2OS cells expressing Fis1TM-mCherry (OMM, grey) and su9-BFP (mitochondrial matrix, red) during membrane-pulling via FluidFM. Yellow triangle indicates position of the aperture inside the cell. 0 s, Δp = 0 mbar. 1 s - 4 s, Δp = -50 mbar. Scale bar: 5 µm (left image), 2 µm (images on the right). (**C**) Time-lapse of mitochondrial network upon pulling. Arrowheads indicate sites of a fission event. Scale bar: 10 µm. See also: Supplementary Movie 5. (**D)** Image series U2OS-cells overexpressing Drp1-mCherry (cyan) and quantified fluorescent signal along mitochondrial tubes during force induced shape transition of pulled mitochondria. (**E**) U2OS cell expressing kinesin-FRP (minus tail) and FKBP-Fis1. Mitochondria fragment following addition of rapamycin. Scale bars: 10 µm.

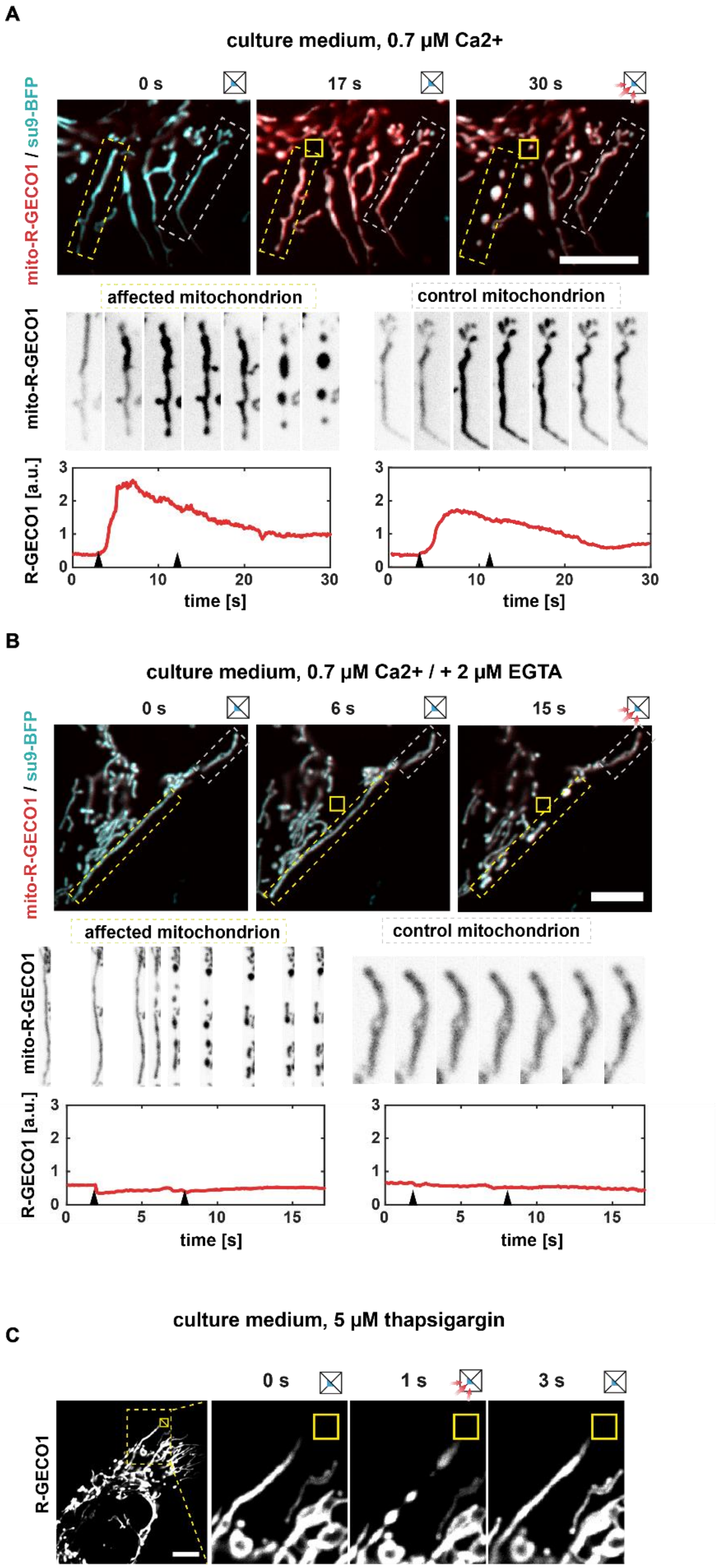
A
culture medium, 0.7 µM Ca2+
0 s
17 s
30 s
mito-R-GECO1 / su9-BFP
affected mitochondrion
control mitochondrion
mito-R-GECO1
R-GECO1 [a.u.]
time [s]
B
culture medium, 0.7 µM Ca2+ / + 2 µM EGTA
0 s
6 s
15 s
mito-R-GECO1 / su9-BFP
affected mitochondrion
control mitochondrion
mito-R-GECO1
R-GECO1 [a.u.]
time [s]
C
culture medium, 5 µM thapsigargin
0 s
1 s
3 s
R-GECO1

**Supplementary Fig. 6.**

Mitochondrial shape transition is $Ca^{2+}$-independent. (**A** and **B**) $Ca^{2+}$ imaging series of mitochondrial shape changes. U2OS cells express su9-BFP (mitochondrial matrix, cyan) and the $Ca^{2+}$ -sensing fluorophore mito-R-GECO1 (red). The upper panel shows an overlay of the two fluorophores during extraction. The middle panel shows an enlarged section of (left panel) a mitochondrion directly adjacent to the cantilever tip and (right panel) a peripheral mitochondrion. The bottom panel shows the total fluorescence intensity of mito-R-GECO1 of the displayed mitochondria during the manipulation process. Yellow boxes indicate position of the cantilever aperture, arrows indicate time point of application of $-\Delta p$. (**A**) Experiment done in culture medium containing 0.7 µM calcium. (**B**) Experiment executed in culture medium after addition of 2 µM EGTA. (**C**) Pulling experiment of an individual mitochondrial tube 30 minutes after the addition of 5 µM thapsigargin. n = 8. Scale bars: 10 µm.

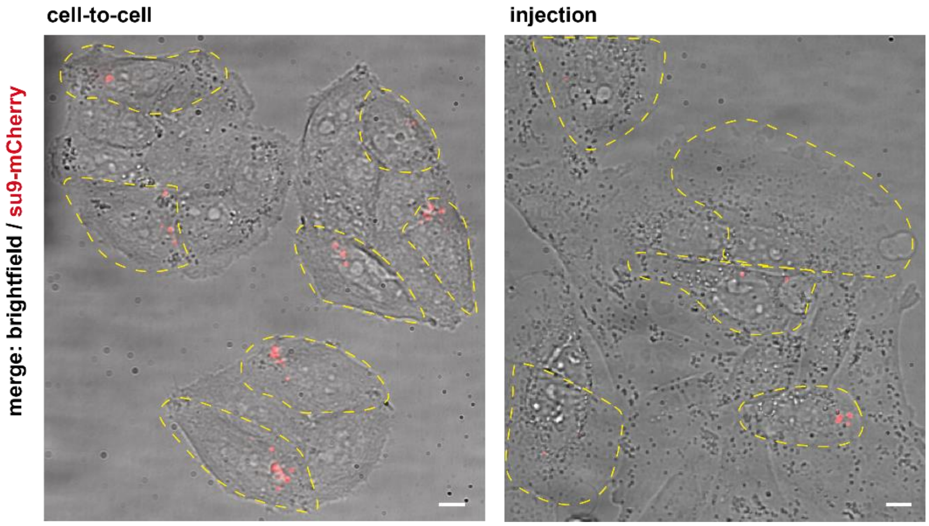

**Supplementary Fig. 7.**
Images of Hela cells post mitochondrial transplantations via the cell-to-cell and the injection approach. Images show and overlay of brightfield (grey) and the transplant (su9-mCherry, red). Transplanted cells are outlined in yellow. Scale bars: 10 µm

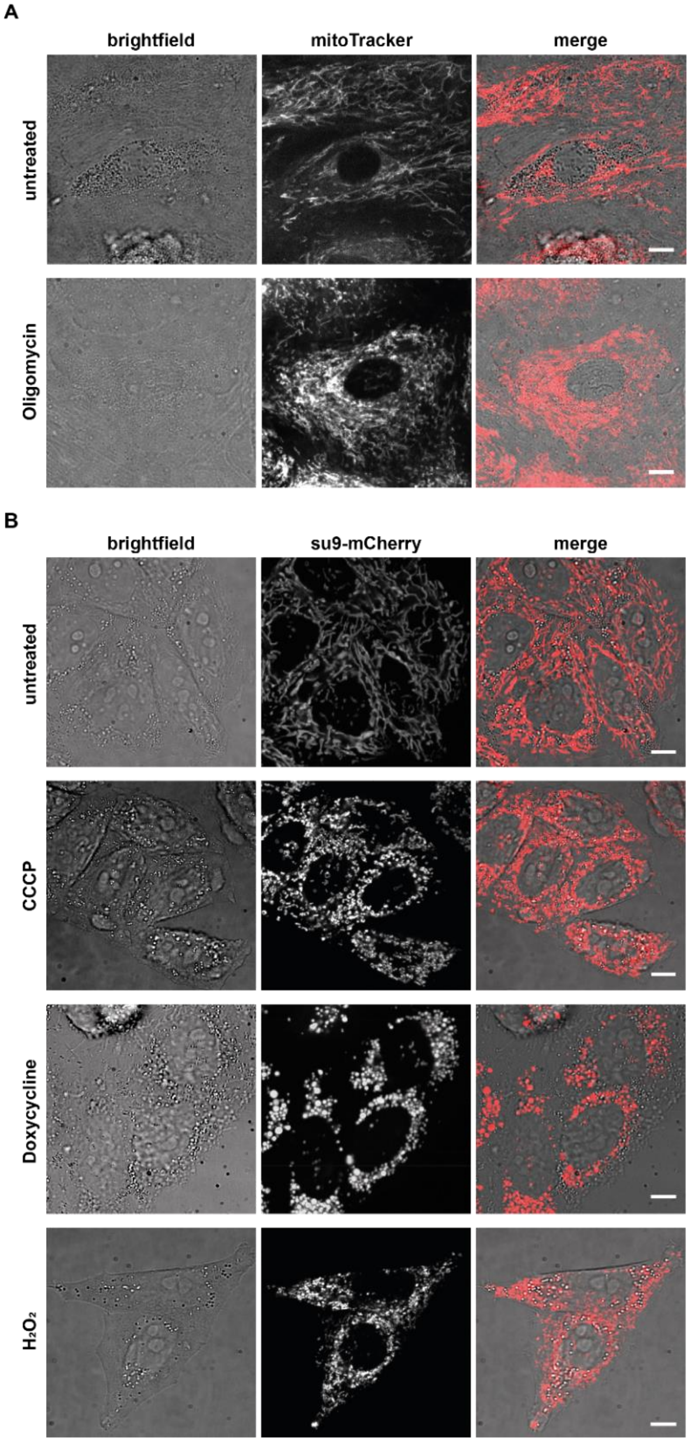

**Supplementary Fig. 8.**
Fusion states of the mitochondrial network of HEKa cells upon drug-treatments. (**A**) HEKa cells in culture medium, the mitochondrial network is visualized using mitoTracker Green. Cells in the lower panel were treated with 4 µM Oligomycin for 22 h. (**B**) Hela cells in culture medium, the mitochondrial matrix is visualized via permanent expression of su9-mCherry. Treatments: 10 µM CCCP for 3 h, 8 µM Doxycycline for 24 h, 750 µM $H_2O_2$ for 3h. Scale bars: 10 µm.

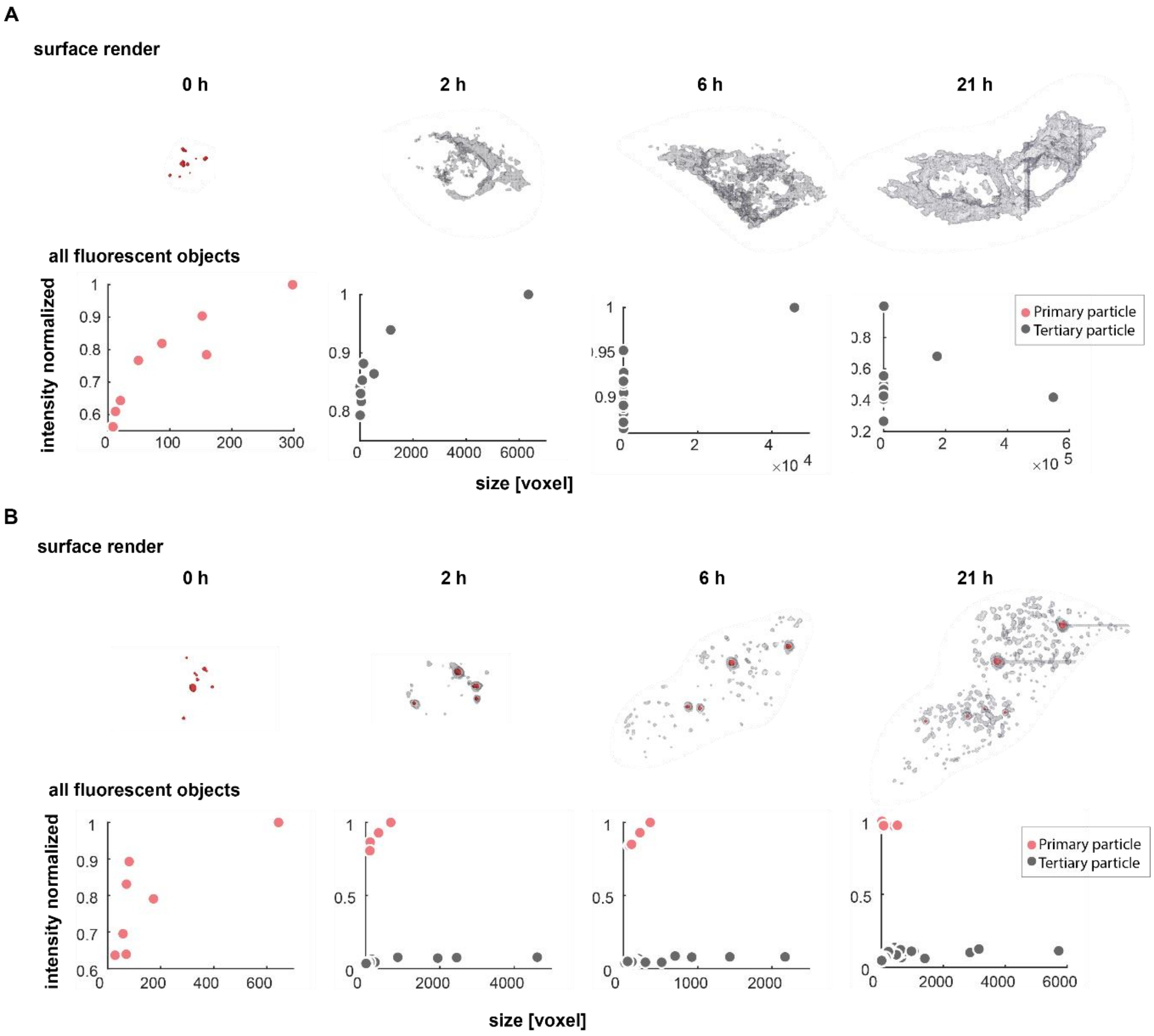

**Supplementary Fig. 9.**

Visualisation and analysis of transplanted mitochondria in single Hela cells. Top: surface render of the total fluorescence of the transplant over time bottom: objects plotted by size in pixel and normalized fluorescence intensity. Objects carrying the high fluorescence intensity of the initial transplant at the 0 h time point are depicted in red; objects with low fluorescence are depicted in grey. (**A**) Hela cell showing mitochondrial acceptance of 8 mitochondria within 2 h. (**B**) Hela cell showing full degradation of the transplant, 7 mitochondria were transplanted.

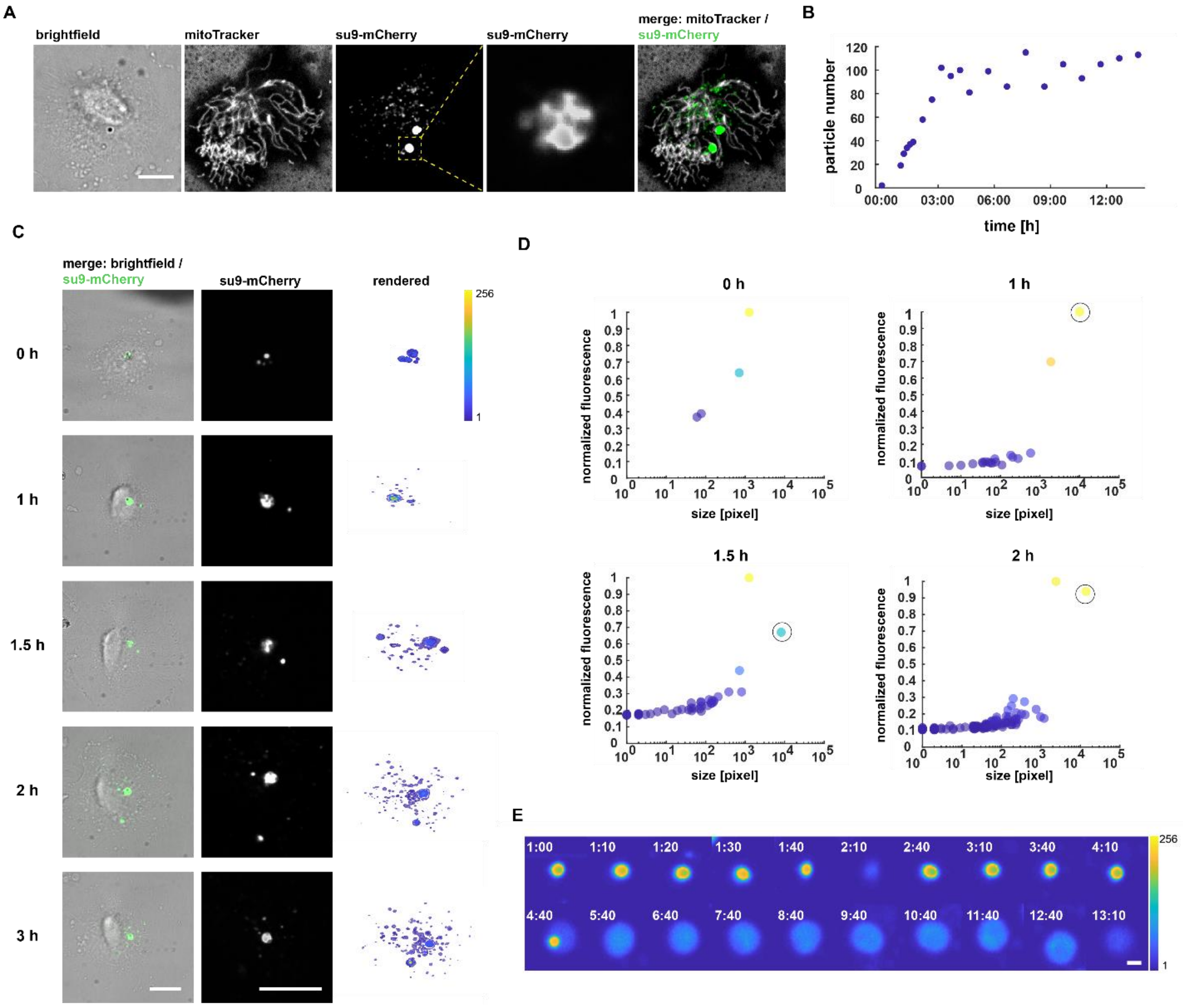

**Supplementary Fig. 10.**
Exemplary quantification of mitochondrial degradation in HEKa cells. (**A**) Fluorescence microscopy images of a cell 19 h post mitochondrial transplantation, 4 mitochondria were initially transplanted. The enlarged section shows a presumptive mitophagosomal structure. Scale bar: 10 µm. (**B**) Total number of fluorescent objects within the host cell over time. (**C**) Time-lapse images of mitochondrial degradation. Surface-rendered images of all detected fluorescent images show the increasing number and spatial distribution of particles over time. Scale bar: 10 µm. (**D**) Scatter plots of objects plotted by size in pixel (logarithmic scale) and normalized fluorescence intensity. Outlined dot shows the largest tertiary particle shown in the central panel in c. **e**, degradation of individual mitochondria over time [h]. After fusion with a presumptive mitophagosomal structure at 4:40 hours, the degradation progresses for over 8 hours. Scale bar: 1 µm.

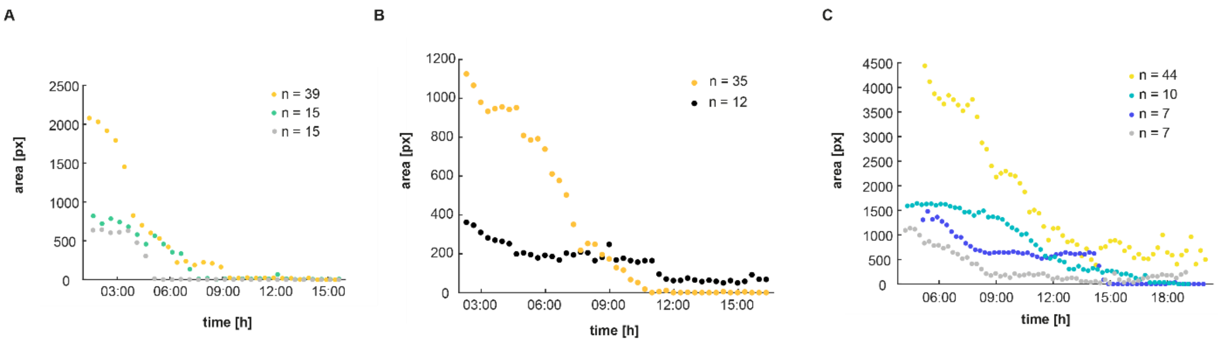

**Supplementary Fig. 11.**
Fluorescent traces of the mitochondrial transplant and directly derived particles in single HEKa cells over time. Plots show the total volume occupied by unfused mitochondrial transplant over time in various conditions. The number of initially transplanted mitochondria per cell is depicted on the upper right. (**A**) Transplant was treated with CCCP and Oligomycin. (**B**) Transplant was treated with Doxycycline, CCCP and Oligomycin. (**C**) Transplant was treated with Doxycycline.

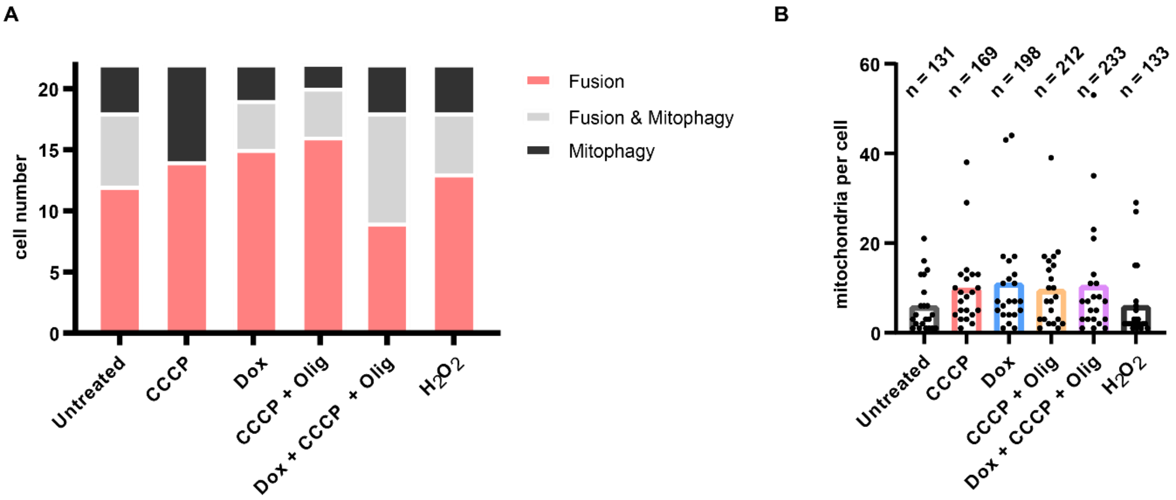

**Supplementary Fig. 12.**
(**A**) Fusion and degradation behavior of drug-compromised mitochondrial transplants in HEKa cells. Each condition was tested with 22 cells. **(B**) Distribution of mitochondrial quantity transplanted per cell from all conditions tested in a (22 cells for each conditions). Line shows median value.

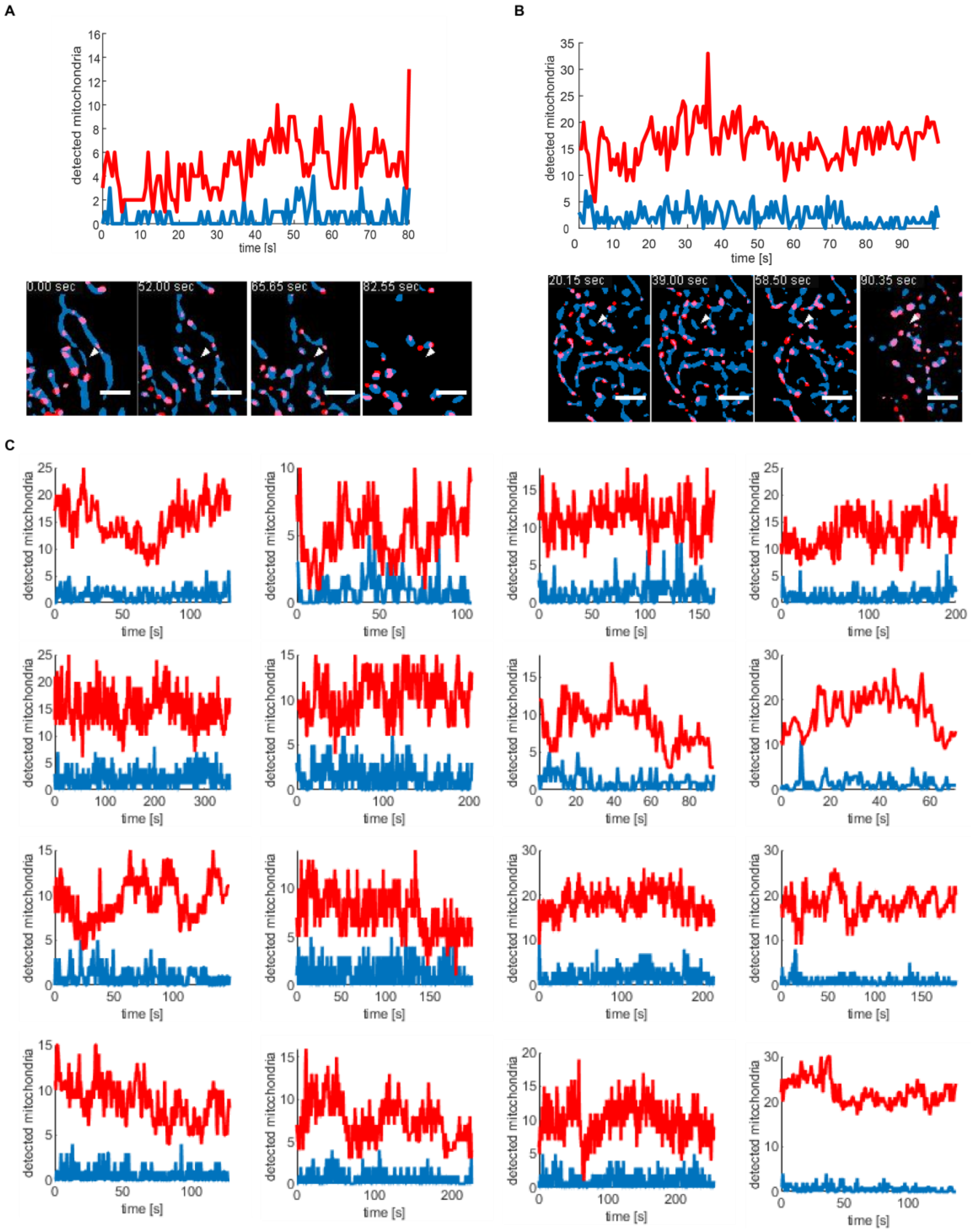

**Supplementary Fig. 13.**
Tracing of p55-nucleoids overlap with mitochondria during extraction via FluidFM. (**A** and **B**) traces of separated mitochondria (su9-BFP) showing overlap with p55-GFP signals in red and of separated mitochondria showing no traces of p55-GFP in blue. White arrowhead indicates cantilever apex position. Panels below show the extraction sites comprising the area used for data acquisition. The mitochondrial network fluorescence is shown in blue, p55-GFP speckles are shown in red. Scale bars: 5 µm. (**C**) Traces of all cells analyzed in this manner.

**Captions for supplementary movies:**

**Supplementary Movie 1. Organelle extraction using FluidFM.** Brightfileld video of HEK cells being extracted with a FluidFM cantilever prefilled with perfluorooctane. Scale bar = 10 µm.

**Supplementary Movie 2. ER-extraction from a COS7 cell using FluidFM**. Panels from left to right: Mitochondrial matrix, su9-BFP; ER, Sec61-GFP; Merge of panels one and two: mitochondria (red) and ER (cyan). Arrowheads mark the site of extraction. Scale bar: 5 µm.

**Supplementary Movie 3. Cell-wide view of ER-extraction from a COS7 cell using FluidFM.** Channels from left to right: Mitochondrial matrix, su9-BFP; ER, Sec61-GFP; merge of channels one and two: mitochondria (red) and ER (cyan). Arrowheads mark the site of extraction. Scale bars: 10 µm.

**Supplementary Movie 4. Extraction of mitochondria from a U2OS cells using FluidFM.** Mitochondrial matrix is labelled via su9-BFP. Arrowhead indicates site of extraction. Left side: cell wide view of the extraction process; scale bar: 10 µm. Right side: enlarged view of the extraction site; scale bar: 5 µm.

**Supplementary Movie 5. Extraction of singular mitochondrial sphere from a U2OS cell using FluidFM.** Mitochondrial matrix is labelled via su9-BFP. Arrowhead indicates site of extraction. Left side: cell wide view of the extraction process; scale bar: 10 µm. Right side: enlarged view of the extraction site; scale bar: 5 µm.

**Supplementary Movie 6. Pearling of mitochondrial tubes upon exertion of pulling force**. Channels from left to right: Mitochondrial matrix: su9-BFP; outer mitochondrial membrane: Fis1TM-mCherry; merge of mitochondrial matrix (red) and outer mitochondrial membrane (cyan). Arrowheads indicate site of extraction. Scale bars: 5 µm.

**Supplementary Movie 7. Calcium imaging during mitochondrial extraction without membrane disruption via FluidFM.** Movie of an individual U2OS cell. Channels from left to right: $Ca^{2+}$ sensor within the mitochondrial matrix, mito-R-GECO1; mitochondrial matrix, su9-BFP; merge of the $Ca^{2+}$ sensor (red) and the mitochondrial matrix label (cyan). Arrowheads indicate site of extraction. Scale bar: 10 µm.

**Supplementary Movie 8. Calcium imaging during membrane disruption via FluidFM.** Movie of the individual U2OS cell shown in Movie S7. Channels from left to right: $Ca^{2+}$ sensor within the mitochondrial matrix, mito-R-GECO1; mitochondrial matrix, su9-BFP; merge of the $Ca^{2+}$ sensor (red) and the mitochondrial matrix label (cyan). Arrowhead indicates site of extraction. Scale bar: 10 µm.

**Supplementary Movie 9. Calcium imaging during extraction with mild membrane disruption.** Channels from left to right: $Ca^{2+}$ sensor within the mitochondrial matrix, mito-R-GECO1; mitochondrial matrix, su9-BFP; merge of the $Ca^{2+}$ sensor (red) and the mitochondrial matrix label (cyan). Arrowheads indicate site of extraction. Scale bars: 10 µm.

**Supplementary Movie 10. Calcium imaging during extraction with mild membrane disruption after addition of EGTA.** Channels from left to right: $Ca^{2+}$ sensor within the mitochondrial matrix, mito-R-GECO1; mitochondrial matrix, su9-BFP; merge of the $Ca^{2+}$ sensor (red) and the mitochondrial matrix label (cyan). Arrowheads indicate site of extraction. Scale bars: 10 µm.

**Supplementary Movie 11. Fusion of a singular transplanted mitochondrial sphere with the mitochondrial network of a U2OS cell.** Channels from left to right: Mitochondrial matrix of the transplanted mitochondrion, su9-mCherry; mitochondrial matrix of the host cell network, su9-BFP; merge: transplant mitochondria (red) and host network (cyan). Scale bar: 10 µm.

**Supplementary Movie 12. Time-lapse images of mitochondrial acceptance in HEKa cells.** Left: Brightfield images. Right: Fluorescence signal of the transplanted mitochondria, su9- 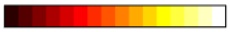 mCherry in the 'hot' colormap, Matlab R2018a: 1 256. The amount of the initially transplanted mitochondria are depicted in the first and last frame next to the respective cell.

**Supplementary tables:**

**Supplementary Table 1. Viability of HeLa cells post mitochondrial transplantation: Injection of purified mitochondria, extracted from bulk cultured cells**

Control - PCR-amplified U2OS mtDNA was mixed with PCR-amplified HeLa mtDNA to the following concentrations: Ctl1 - 0.1 % / Ctl2 - 0.5 % / Ctl3 - 1 %

| **position** | **Ctl1.A** | **Ctl1.C** | **Ctl1.G** | **Ctl1.T** | **Total reads** | **HeLa mtDNA [%]** | **U2OS mtDNA [%]** | **average [%]** |
|---|---|---|---|---|---|---|---|---|
| 15959 | 3 | 0 | 20059 | 18 | 20080 | 99.90 | 0.09 | 0.10 |
| 16069 | 0 | 19831 | 0 | 20 | 19851 | 99.90 | 0.10 | |
| 16108 | 0 | 19650 | 0 | 20 | 19670 | 99.90 | 0.10 | |
| 16126 | 0 | 19 | 0 | 19639 | 19658 | 99.90 | 0.10 | |
| **position** | **Ctl2.A** | **Ctl2.C** | **Ctl2.G** | **Ctl2.T** | **Total reads** | **HeLa mtDNA [%]** | **U2OS mtDNA [%]** | **average [%]** |
| 15959 | 5 | 0 | 28675 | 120 | 28800 | 99.57 | 0.42 | 0.47 |
| 16069 | 1 | 28388 | 0 | 141 | 28530 | 99.50 | 0.49 | |
| 16108 | 0 | 28139 | 0 | 140 | 28279 | 99.50 | 0.50 | |
| 16126 | 0 | 137 | 0 | 28129 | 28266 | 99.52 | 0.48 | |
| **position** | **Ctl3.A** | **Ctl3.C** | **Ctl3.G** | **Ctl3.T** | **Total reads** | **HeLa mtDNA [%]** | **U2OS mtDNA [%]** | **average [%]** |
| 15959 | 2 | 0 | 25492 | 164 | 25658 | 99.35 | 0.64 | 0.77 |
| 16069 | 1 | 25226 | 0 | 210 | 25437 | 99.17 | 0.83 | |
| 16108 | 1 | 25021 | 0 | 210 | 25232 | 99.16 | 0.83 | |
| 16126 | 0 | 201 | 0 | 25016 | 25217 | 99.20 | 0.80 | |

Transplanted - biological replicates 1-3 (i)

| position | TP1. A | TP1. C | TP1. G | TP1. T | Total reads | HeLa mtDNA [%] | U2OS mtDNA [%] | average [%] |
|---|---|---|---|---|---|---|---|---|
| 15959 | 0 | 0 | 27638 | 658 | 28296 | 97.67 | 2.33 | 2.58 |
| 16069 | 0 | 27360 | 0 | 757 | 28117 | 97.31 | 2.69 | |
| 16108 | 0 | 27296 | 0 | 750 | 28046 | 97.33 | 2.67 | |
| 16126 | 0 | 734 | 0 | 27263 | 27997 | 97.38 | 2.62 | |
| **position** | **TP2. A** | **TP2. C** | **TP2. G** | **TP2. T** | **Total reads** | **HeLa mtDNA [%]** | **U2OS mtDNA [%]** | **average [%]** |
| 15959 | 1 | 0 | 32063 | 498 | 32562 | 98.47 | 1.53 | 1.70 |
| 16069 | 1 | 31738 | 0 | 571 | 32310 | 98.23 | 1.77 | |
| 16108 | 0 | 31674 | 0 | 571 | 32245 | 98.23 | 1.77 | |
| 16126 | 0 | 560 | 0 | 31662 | 32222 | 98.26 | 1.74 | |
| **position** | **TP3. A** | **TP3. C** | **TP3. G** | **TP3. T** | **Total reads** | **HeLa mtDNA [%]** | **U2OS mtDNA [%]** | **average [%]** |
| 15959 | 0 | 0 | 15191 | 221 | 15412 | 98.57 | 1.43 | 1.58 |
| 16069 | 0 | 15097 | 0 | 253 | 15350 | 98.35 | 1.65 | |
| 16108 | 1 | 15081 | 0 | 252 | 15334 | 98.35 | 1.64 | |
| 16126 | 0 | 247 | 0 | 15084 | 15331 | 98.39 | 1.61 | |

Injected mitochondria from bulk isolation - biological replicates 1-3 (ii)

| **position** | **Inj1.A** | **Inj1.C** | **Inj1.G** | **Inj1.T** | **Total reads** | **HeLa mtDNA [%]** | **U2OS mtDNA [%]** | **average [%]** |
|---|---|---|---|---|---|---|---|---|
| 15959 | 0 | 0 | 12555 | 79 | 12634 | 99.37 | 0.63 | 0.70 |
| 16069 | 0 | 12434 | 0 | 92 | 12526 | 99.27 | 0.73 | |
| 16108 | 2 | 12403 | 0 | 92 | 12497 | 99.25 | 0.74 | |
| 16126 | 1 | 88 | 0 | 12398 | 12487 | 99.29 | 0.70 | |
| **position** | **Inj2.A** | **Inj2.C** | **Inj2.G** | **Inj2.T** | **Total reads** | **HeLa mtDNA [%]** | **U2OS mtDNA [%]** | **average [%]** |
| 15959 | 0 | 0 | 23824 | 116 | 23940 | 99.52 | 0.48 | 0.56 |
| 16069 | 0 | 23609 | 0 | 137 | 23746 | 99.42 | 0.58 | |
| 16108 | 1 | 23567 | 1 | 140 | 23709 | 99.40 | 0.59 | |
| 16126 | 0 | 135 | 0 | 23569 | 23704 | 99.43 | 0.57 | |
| **position** | **Inj3.A** | **Inj3.C** | **Inj3.G** | **Inj3.T** | **Total reads** | **HeLa mtDNA [%]** | **U2OS mtDNA [%]** | **average [%]** |
| 15959 | 0 | 0 | 29343 | 35 | 29378 | 99.88 | 0.12 | 0.14 |
| 16069 | 0 | 29123 | 0 | 44 | 29167 | 99.85 | 0.15 | |
| 16108 | 2 | 29080 | 0 | 45 | 29127 | 99.84 | 0.15 | |
| 16126 | 0 | 42 | 0 | 29075 | 29117 | 99.86 | 0.14 | |

Mixed - biological replicates 1-3 (iii)

| **position** | **Mixed1.A** | **Mixed 1.C** | **Mixed.G** | **Mixed1.T** | **Total reads** | **HeLa mtDNA [%]** | **U2OS mtDNA [%]** | **average [%]** |
|---|---|---|---|---|---|---|---|---|
| 15959 | 2 | 0 | 38482 | 3 | 38487 | 99.99 | 0.01 | 0.00 |
| 16069 | 0 | 38329 | 0 | 1 | 38330 | 100.00 | 0.00 | |
| 16108 | 0 | 38281 | 0 | 0 | 38281 | 100.00 | 0.00 | |
| 16126 | 1 | 0 | 0 | 38265 | 38266 | 100.00 | 0.00 | |
| **position** | **Mixed 2.A** | **Mixed 2.C** | **Mixed.G** | **Mixed2.T** | **Total reads** | **HeLa mtDNA [%]** | **U2OS mtDNA [%]** | **average [%]** |
| 15959 | 2 | 0 | 32911 | 0 | 32913 | 99.99 | 0.00 | 0.00 |
| 16069 | 0 | 32764 | 0 | 1 | 32765 | 100.00 | 0.00 | |
| 16108 | 1 | 32725 | 0 | 1 | 32727 | 99.99 | 0.00 | |
| 16126 | 0 | 0 | 0 | 32719 | 32719 | 100.00 | 0.00 | |
| **position** | **Mixed 3.A** | **Mixed 3.C** | **Mixed3.G** | **Mixed3.T** | **Total reads** | **HeLa mtDNA [%]** | **U2OS mtDNA [%]** | **average [%]** |
| 15959 | 3 | 0 | 38647 | 0 | 38650 | 99.99 | 0.00 | 0.00 |
| 16069 | 0 | 38446 | 0 | 0 | 38446 | 100.00 | 0.00 | |
| 16108 | 0 | 38402 | 0 | 1 | 38403 | 100.00 | 0.00 | |
| 16126 | 0 | 0 | 0 | 38389 | 38389 | 100.00 | 0.00 | |

**Supplementary Table 2. Primers used in this study**

| Name | Binding site mtDNA | Sequence 5' – 3' |
| --- | --- | --- |
| Primer 1 | 15720 | ATTGACTCCTAGCCGCAGAC |
| Primer 2 | 16298 | AAGGGTGGGTAGGTTTGTTG |
